# IL4-driven microglia modulate stress resilience through BDNF-dependent neurogenesis

**DOI:** 10.1101/2020.02.01.929646

**Authors:** Jinqiang Zhang, Peijing Rong, Lijuan Zhang, Hui He, Tao Zhou, Yonghua Fan, Li Mo, Qiuying Zhao, Yue Han, Shaoyuan Li, Yifei Wang, Wan Yan, Huafu Chen, Zili You

## Abstract

Adult neurogenesis in the dentate gyrus of the hippocampus is regulated by specific groups of microglia and is functionally implicated in behavioral responses to stress. However, the role of microglia in modulating hippocampal neurogenesis in stress responses remains poorly understood. Here we investigated the effects of IL4-driven Arg1^+^ microglia in the restoration of hippocampal neurogenesis and conferment of stress resilience. We found that low IL4 levels in the hippocampus of mice was associated with greater stress vulnerability and, conversely, overexpression of IL4 in the hippocampus induced a large number of Arg1^+^ microglia and ameliorated stress-induced depressive-like behaviors. Knockdown of microglial IL4 receptors in the hippocampus of mice exacerbated the stress-induced inflammatory response and abolished the antidepressant effects of IL4 overexpression. Enhancement or inhibition of IL4 signaling in hippocampal microglia modulated neurogenesis, and blockade of neurogenesis abolished the resilience to stress-induced depression. We further show that IL4-activated microglia is associated with upregulation of BDNF levels and neurogenesis. Taken together, our findings suggest that IL4-driven microglia in the hippocampus trigger BDNF-dependent neurogenesis in response to chronic stress, helping protect against depressive-like symptoms. These findings identify the modulation of a specific microglial phenotype as a treatment strategy for mood disorders.

**In Brief:** Zhang et al. show that IL4-induced Arg1^+^ microglia restore hippocampal neurogenesis and promote resilience against stress in mice by increasing BDNF levels. Targeting microglia with immunomodulatory factors may be a strategy for treating mood disorders.

**Highlights:**

1. Vulnerability to stress in mice is associated with reduced IL4 signaling in the hippocampus
2. Brain-derived IL4 promotes adult hippocampal neurogenesis and stress resistance by driving Arg1^+^ microglia
3. IL4-driven Arg1^+^ microglia enhance hippocampal neurogenesis via a BDNF-dependent pathway

## Introduction

It is well-known that prolonged stress can induce cascading effects in the inflammatory system and lead to neuropsychiatric disorders, such as depression and anxiety, which can affect neurogenesis ^1^. New neurons are generated throughout adulthood in two regions of the brain, the dentate gyrus (DG) in the hippocampus and the subventricular zone ^2^. Neural stem/progenitor cells (NSPCs) undergo proliferation, differentiation, survival and maturation into new neurons, which eventually integrate into the neural network ^3^. These processes are regulated negatively by stressful experiences and positively by treatment with antidepressant drugs through altering the neurogenic microenvironment ^2^. These adult-born neurons are critical for mood control and behavioral output ^2, 4^. Alterations in the magnitude of this continuous production of new neurons are associated with both the pathophysiology of depression and the efficacy of antidepressants ^1, 5, 6^.

Microglia, the principal immune cells in the brain, play central roles in immune surveillance and maintenance of brain homeostasis ^7^. Studies have shown that microglia can regulate neurogenesis by secreting various factors that modulate the neurogenic microenvironment ^8, 9^. Under stress or pathological conditions, microglia secrete inflammatory mediators that disrupt neuronal function and impair neurogenesis, increasing the vulnerability to stress and promoting the occurrence and development of depression ^9–11^. On the other hand, recent evidence showed that microglial ‘alternative activation’ leads to an increase in adult neurogenesis in response to exercise and environmental enrichment, which contribute to normal behavior ^12, 13^. The role of microglia in neurogenesis depends on the balance between the soluble factors released by microglial cells under different activation scenarios. Microglial brain-derived neurotrophic factor (BDNF) is an important mediator of microglia-to-neuron communication, and contributes to many facets of brain function via its high-affinity receptor, tropomyosin-related kinase B (TrkB) ^14, 15^. Microglial BDNF may also be involved in neurobehavioral plasticity and neurogenesis, and therefore may underlie the pathophysiology of depression ^12, 16^.

Interleukin 4 (IL4) is a multifunctional cytokine expressed in the brain and is involved in the regulation of inflammatory responses and physiological processes of the central nervous system (CNS) ^17–19^. IL4 deficiency weakens the resilience against stress-induced depression, while increasing IL4 levels can alleviate depressive-like behaviors ^20–23^. Studies have shown that IL4 can reprogram microglia toward an Arg1^+^ phenotype for maintaining brain homeostasis, neuroprotection and tissue repair ^24–27^. However, the effects of IL4-induced Arg1^+^ microglia on neurogenesis and stress resilience are poorly understood.

In light of the role of microglia in the pathogenesis of depressive disorders, we hypothesized that promoting a neuroprotective phenotype of microglia would improve stress responses and depressive behaviors. Consistent with this expectation, we found that in mice exposed to chronic mild stress (CMS), overexpression of hippocampal IL4 drove Arg1^+^ microglia to promote neurogenesis, which in turn attenuated depressive-like behaviors.

## Material and methods

### Animals

*CX_3_CR1^Cre/ER^* mice on a C57BL/6J background were purchased from The Jackson Laboratory. Wild-type (WT) C57BL/6J mice were purchased from the Laboratory Animal Center of Sichuan Academy of Medical Sciences (Chengdu, China). All mice were housed under standard conditions with free access to food and water. All animal experiments were approved by the Ethics Committee of the University of Electronic Science and Technology of China and carried out in strict accordance with the US National Institutes of Health *Guide for the Care and Use of Laboratory Animals* (8th edition, revised 2010).

### Chronic Mild Stress (CMS)

Male mice aged 8–10 weeks at the start of experiments were caged individually and subjected to a CMS protocol ^10^. To establish an animal model of depression, the CMS paradigm was conducted over a 4-week period and consisted of daily exposure to three stressors in random order. To screen the vulnerable animals, the CMS paradigm was conducted over a 3-week period. These stressors included empty water bottles (12 h), food deprivation (12 h), tail clipping (10 min), restraint (2 h), lights-off for 3 h during the daylight phase, cage shaking (1 h), cage tilting (45°, 24 h), reversal of the light–dark cycle (24 h), strobe lighting (12 h), damp bedding (24 h) and a soiled cage (24 h).

### Animal screening

Mice were subjected to CMS for 3 weeks, then subdivided as described ^28^ into groups of high-susceptible (HS) or low-susceptible (LS) animals. The mice were considered LS if their immobility time in the forced swim test and tail suspension test, and their sucrose intake (see below) were within one standard deviation of the mean values for control mice. If these values fell outside this threshold, animals were considered HS.

### Expression systems and stereotactic injection

Recombinant adeno-associated virus (rAAV) 2/9 was produced by transfecting 293T cells with three plasmids: an AAV vector expressing both IL4 and enhanced green fluorescent protein (EGFP), IL4 alone or EGFP alone; an AAV helper plasmid (pAAV Helper); and an AAV Rep/Cap expression plasmid. At 72 h after transfection, cells were collected and lysed using a freeze-thaw procedure. AAV-IL4 or AAV were diluted to 1 × 10^10^ transforming units (TU)/mL and injected (0.5 µL) bilaterally in the hippocampus (Bregma: 2.3 mm, L: 1.8 mm, V: 2.0 mm).

To downregulate expression of the IL4 receptor α-chain (IL4Rα) in hippocampal microglia, a lentivirus encoding a short hairpin RNA (shRNA) against the receptor was constructed as described ^29^. The lentivirus contained the cassette u6-LoxP-stop-LoxP-shRNA-IL4Rα. To construct shRNA-IL4Rα, oligonucleotides containing antisense sequences were connected with a hairpin loop, followed by a poly termination signal. The sequences targeting IL4Rα (GenBank accession: NM_001008700, ID: 16190) in this experiment were 5′-GCG CTG TAT GGA GCT GTT TGA-3′ and 5′-TCA AAC AGC TCC ATA CAG CGC-3′.

For *in vivo* viral injections, viral vectors were bilaterally targeted to the hippocampus (Bregma: 2.3 mm, L: 1.8 mm, V: 2.0 mm). Animals received intracranial viral injections while under 10% pentobarbital anesthesia using a Kopf stereotactic apparatus. Mice were secured using ear bars and a head holder. A Micro-1 microsyringe pump controller (RWD life science) was used to inject viral vectors. A 33-GA needle was lowered 2 mm over 5 min resulting in delivery of 0.25 μL viral vectors (2.5 × 10^9^ TU) into each hippocampus. After injection, the syringe was held in place for 5 min to avoid back-flow. Animals were sutured and placed on a heating pad for recovery. Mice behavior was analyzed at 2 weeks after these injections (see below). Tamoxifen (Sigma) was dissolved in warm sunflower seed oil at a concentration of 40 mg/ml and injected intraperitoneally at 100 mg/kg for 7 days to induce Cre recombinase expression.

### Behavioral analyses

#### Sucrose preference test (SPT)

The sucrose preference test was performed as described ^10^. Mice were individually housed, deprived of food and water for 12 h, and then given access to 1% sucrose solution (A) and water (B) for 2 h. The bottle positions were switched daily to avoid a side bias. The sucrose preference was calculated each week for each mouse using the formula: 100 × [VolA / (VolA + VolB)]. The sucrose consumption was normalized to body weight for each mouse.

#### Locomotor activity test (LAT)

Each mouse was placed in an autonomous movement instrument with a 36-point infrared ray passive sensor system (ZZ-6, Taimeng Tech, Chengdu, China). All mice were allowed to habituate to the environment for 1 min before the test. The total locomotor activity (standing and movement) was automatically recorded for each mouse during 10 min.

#### Forced swim test (FST)

Each mouse was placed for 15 min in a cylinder containing 15 cm of water. After 24 h, the animals were placed again in the cylinder for 6 min. The duration of immobility (including staying afloat and movements to stay afloat) was measured using FST100 software (Taimeng Tech) during the last 4 min of swimming time.

#### Tail suspension test (TST)

The TST was performed as described ^10^. Mice were elevated by securing the tail 30 cm above the ground with adhesive plaster. Mice were isolated from one another using a black cardboard. Mice were recorded for 6 min, and time spent immobile during that period was determined by observers blind to mouse group allocations.

#### Coat score assay

The physical state of the fur was evaluated weekly in the coat score assay as described ^30^. The total coat score was measured as the sum of the scores for seven different body parts: head, neck, forepaws, dorsal coat, ventral coat, hindpaws and tail. For each area, a score of 1 was given for a well-groomed coat and a score of 0 was given for an unkempt coat.

#### Novelty-suppressed feeding test (NSFT)

Mice were deprived of food and water for 12 h before the test, and then each mouse was placed for 5 min in a rectangular chamber (40 × 40 × 30 cm) containing a sugar pill in the center of the chamber. The time it took for a mouse to pick up the sugar with its forelimb was recorded as latency using a camera system.

#### Open field test (OFT)

The mice were gently placed in the center of a rectangular chamber (40 × 40 × 30 cm) and left to explore the area for 5 min. The digitized image of the path taken by each mouse was stored, and the locomotion activity and time spent in the center were analyzed post hoc for each mouse using OFT100 software (Taimeng Tech).

#### Gene expression analysis

Total RNA was isolated from mouse hippocampus using Isol-RNA Lysis Reagent (5 Prime). Quantitative real-time PCR was performed, and gene expression was quantified using the -ΔΔCt method. Primer sequences are listed in Table S1.

#### Enzyme-linked immunosorbent assay (ELISA)

Hippocampal sections of mice were sonicated in RIPA-buffer containing protease inhibitors. Cytokine concentrations were quantified using ELISA kits (QuantiCyto, China) according to the manufacturer’s protocol. Detection limits were 8 pg/mL for IL4 and Arg1, and 4 pg/mL for iNOS, TGFß and BDNF.

#### Western blotting

Hippocampal tissue or isolated hippocampal microglia were sonicated in RIPA-buffer containing protease inhibitors. Protein samples were run on 12% Tris-glycine SDS-PAGE gels, transferred to PVDF membrane (0.2 or 0.45 μm), and blotted with antibodies (Abcam) against IL4 (1:1000), IL4Rα (1:1000), STAT6 (1:800), pSTAT6 (1:800) Arg1 (1:1000), TrkB (1:1000) and β-actin (1:20,000). Primary antibody was incubated overnight at 4°C, and secondary antibodies (1:10,000, Abcam) were incubated for 2 h at room temperature. Signals were developed using the ECL-Plus kit (Millipore, USA). Densitometry was performed to quantify signal intensity using ImageJ software (version 1.45 J; National Institutes of Health, Bethesda, MD, USA).

#### BrdU incorporation

To determine cell proliferation in the brain, mice received two intraperitoneal injections of BrdU (50 mg/kg) 8 h apart. To examine progenitor proliferation, mice were sacrificed 24 h after the second injection; to examine progenitor differentiation, mice were sacrificed 1 week after BrdU injection. For examination of neuronal survival in the granular layer, animals were injected with a double dose of BrdU and sacrificed 4 weeks after injection.

#### Tissue preparation and immunohistochemistry

Tissue preparation and staining were performed as described ^10^ using the antibodies listed in Table S2.

#### Isolation of microglia from hippocampus

Hippocampus was isolated from the brain and homogenized into single-cell suspensions, from which microglia were isolated on a Percoll density gradient as described ^31^. In brief, the hippocampus was homogenized, and cell pellets were resuspended in 70% isotonic Percoll. A discontinuous Percoll density gradient (70%, 50%, 35%, 0%) was layered and centrifuged for 20 minutes at 2000g. Enriched microglia were collected from the interphase between 70% and 50% Percoll. Of the cells recovered from this Percoll interphase, approximately 90% of the cells were CD45^int^-CD11b^+^ microglia.

#### Flow Cytometry

Cells were assayed for surface antigens by flow cytometry as previously described ^32^. These microglia were labeled with FITC anti-mouse CD45 (2 µg/mL, Biolegend, USA), APC anti-mouse CD11b (5 µg/mL, Biolegend, USA), APC/Cyanine7 anti-mouse CD86 (1 µg/mL, Biolegend, USA), and PerCP/Cyanine 5.5 CD206 (5 ug/mL, Biolegend, USA) antibodies for 30 min at room temperature. A four-laser Becton-Dickinson FACS Calibur (BD Biosciences) was used to collect the data, and FlowJo software was used for analysis.

#### RNA sequencing

Total RNA was extracted from hippocampal tissue using TRIzol® Reagent according to the manufacturer’s instructions (Invitrogen) and genomic DNA was removed using DNase I (TaKara). Then RNA quality was determined by 2100 Bioanalyser (Agilent) and quantified using the ND-2000 (NanoDrop Technologies). Only high-quality RNA samples (OD260/280 = 1.8 ∼ 2.2, OD260/230 ≥ 2.0, RIN ≥ 6.5, 28S:18S ≥ 1.0, > 2 μg) were used to construct the sequencing library.

RNA purification, reverse transcription, library construction and sequencing were performed according to the manufacturer’s instructions (Illumina, San Diego, CA). The RNA-seq transcriptome library was prepared following TruSeqTM RNA sample preparation Kit from Illumina (San Diego, CA) using 1μg of total RNA. Briefly, mRNA was isolated according to poly A selection method by oligo (dT) beads and then fragmented using fragmentation buffer. Double-stranded cDNA was synthesized using a SuperScript double-stranded cDNA synthesis kit (Invitrogen, CA) with random hexamer primers (Illumina). Then the synthesized cDNA was subjected to end-repair, phosphorylation and ‘A’ base addition according to Illumina’s library construction protocol. The size of libraries was selected for cDNA target fragments of 200–300 bp on 2% Low Range Ultra Agarose followed by PCR amplified using Phusion DNA polymerase (NEB) for 15 cycles. After quantification by TBS380, paired-end RNA-seq library was sequenced on the Illumina NovaSeq 6000 (2 × 150bp read length). All RNA-Seq raw data were uploaded in Sequence Read Archive (SRA: PRJNA602066).

#### Analysis of RNA sequencing data

RNA-Seq data were initially filtered to obtain clean data, including removing reads with adaptors, reads with more than 10% unknown bases or low quality reads (the percentage of low quality bases is over 50% in the read). To identify differential expression genes (DEGs) between two different samples, the expression level of each transcript was calculated according to the fragments per kilobase of exon per million mapped reads (FPKM) method. The package software Empirical analysis of Digital Gene Expression in R (EdgeR) was utilized for differential expression analysis. In addition, functional-enrichment analysis including GO and KEGG was performed to identify which DEGs were significantly enriched in GO terms and metabolic pathways at Bonferroni-corrected P-value ≤ 0.05 compared with the whole-transcriptome background. GO functional enrichment and KEGG pathway analysis were carried out by Goatools.

#### Cell culture and treatments

Neural stem/progenitor cells (NSPCs) and primary microglia were cultured as described ^10^. Microglia were treated with IL4 (20 ng/mL) or IFNγ (10 ng/mL) for 24 h to induce microglial activation ^24^. To assay the short-term effects of microglia on NSPCs, NSPCs were co-cultured with microglia in transwell plates for 24 h in experiments to examine proliferation, or for 3 days in experiments to examine differentiation. To assay the long-term effects of microglia on NSPCs, NSPCs were cultured in microglia-conditioned medium (M-CM) for 24 h in experiments to examine proliferation, or for 7 days in experiments to examine differentiation. The BDNF pathway was blocked using anti-BDNF antibody (1 μg/mL) or BDNF receptor antagonist K252a (100 ng/mL).

#### Imaging and statistical analyses

Images were acquired using a Zeiss AxioImager Z1, and cells were counted manually. Areas of Iba1 or GFAP staining and relative fluorescence intensity were analyzed using Image J software. Branches of microglia were measured using Image-Pro Plus 6.0 (Media Cybernetics, USA). To quantify total cell populations in the hippocampus (microglia, astrocytes, BrdU^+^ cells, BrdU^+^-DCX^+^ cells and BrdU^+^-NeuN^+^ cells), every sixth section (25 μm thick) of the brain containing hippocampus was selected and immunostained. Total numbers of positive cells in all slices per animal were multiplied by six to estimate the number of cells per hippocampus. For measurement of the volume of hippocampus, DG and granule cell layer (GCL), every sixth section (25 μm thick) of the brain containing hippocampus was selected and labelled with DAPI. Total area of hippocampus, DG or GCL in all slices per animal was multiplied by six to estimate the volume of hippocampus, DG and GCL.

All quantitative results were expressed as mean ± SEM. Data were plotted and analyzed statistically using GraphPad Prism 7.0. Potential differences between the mean values were evaluated using one- or two-way analysis of variance (ANOVA), followed by the least significant difference (LSD) test for post hoc comparisons assuming equal variances. Independent-samples *t* tests were used to compare the differences between two groups, unless otherwise specified. Asterisks were used to indicate significance: *p < 0.05, **p < 0.01, and ***p < 0.001. Values > 0.05 were considered not significant (n.s.).

## Results

### Vulnerability to stress is associated with reduced IL4 signaling in the hippocampus

Following exposure to CMS for 3 weeks, C57BL/6J mice were separated into HS and LS subpopulations according to their sucrose preference and immobility time in TST and FST (Fig. 1a). These subpopulations were confirmed by open field test, body weight and coat score (Fig. S1). We examined cytokines, chemokines and neurotrophic factors related to neuroinflammatory processes in the hippocampus and prefrontal cortex, and found that levels of IL4 mRNA in the hippocampus were elevated in LS mice but reduced in HS mice, compared to the control mice (Fig. 1b). In particular, the change in IL4 protein levels in the hippocampus was the most significant outcome (Fig. 1c). Western blotting showed that hippocampal IL4-STAT6 signaling was increased in LS mice, but decreased in HS mice (Fig. 1d). Correlation analysis showed that the IL4 concentration in the hippocampus positively correlated with sucrose preference and negatively correlated with immobility time in TST (Fig. 1e), suggesting that the vulnerability of mice to CMS was associated with reduced IL4 signaling in the hippocampus.

**Fig. 1.**
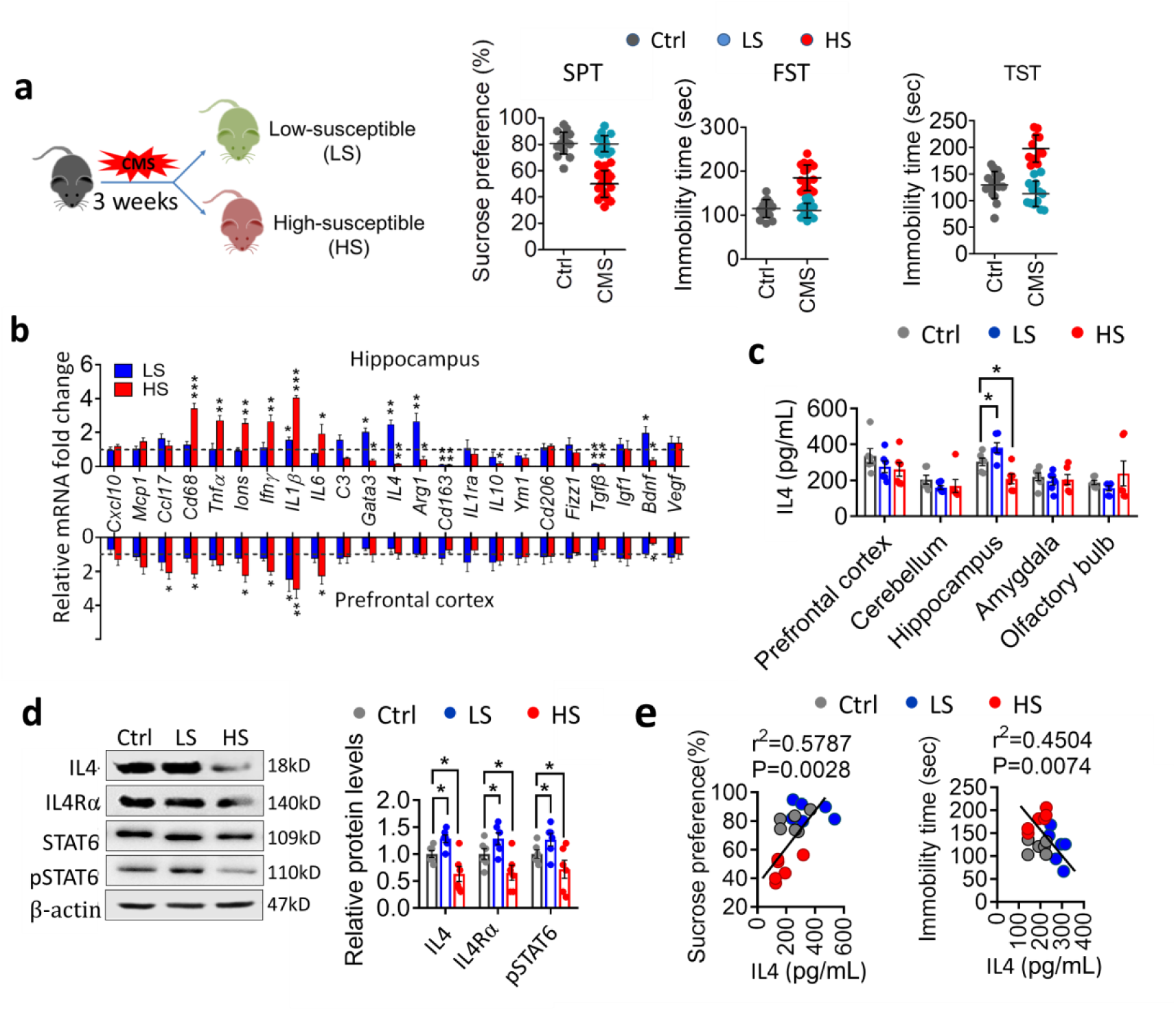
Hippocampal IL4 is involved in resistance of mice to stress. **a**, Behavioral screening of low-susceptible (LS) and high-susceptible (HS) subpopulations of mice exposed to 3-week chronic mild stress (CMS). Results are shown for control mice (n = 20), HS mice (n = 13), and LS mice (n = 11). Data are mean ± SEM. **b**, Quantitative PCR to examine mRNA expression of cytokines, chemokines, growth factors and trophic factors in the hippocampus and prefrontal cortex of HS and LS mice. Data are standardized to control group, and show mean± SEM (n = 4–6 per group). * P < 0.05, ** P < 0.01, *** P < 0.005 *vs* control group (one-way ANOVA with LSD test). **c**, ELISA to assay the protein level of IL4 in prefrontal cortex, cerebellum, hippocampus, amygdala and olfactory bulb of control mice, HS and LS mice. Each circle represents one mouse (n = 6 per group). Data are mean ± SEM. * P < 0.05 (one-way ANOVA with LSD test). **d**, Western blotting shows the levels of IL4, IL4Rα, STAT6 and pSTAT6 in the hippocampus of control mice, HS and LS mice. IL4 and IL4Rα are normalized to β-actin, the pSTAT6 is normalized to STAT6. Data are mean ± SEM (n = 6 per group). * P < 0.05 (one-way ANOVA with LSD test). **e**, Correlation between sucrose preference or immobility duration in TST, and hippocampal IL4 levels. Each circle represents one mouse (n = 6 per group).

### Overexpression of IL4 in the hippocampus induced microglial Arg1^+^ phenotype

Since IL4 is a rapidly degraded glycoprotein of about 20 kDa that cannot cross the blood-brain barrier ^33^, we specifically increased IL4 levels in the hippocampus using rAAV vectors combined with Ef1α promoter encoding murine IL4 to investigate the effect of brain-derived IL4 on depressive-like behaviors. The AAV2/9-IL4-EGFP is more highly infectious to cultured neurons than to astrocytes and microglia (Fig. S2). AAV2/9-IL4-EGFP was stereotactically injected into the hippocampus of C57BL/6J mice, such that EGFP would be co-expressed with IL4. EGFP was mainly localized in NeuN^+^ cells of the hippocampus (Fig. 2a). Another construct encoding murine IL4 without EGFP (AAV-IL4) was stereotactically injected into the hippocampus; in this case, IL4 expression in the hippocampus was confirmed by quantitative PCR, ELISA, and IL4Rα, STAT6 and pSTAT6 were examined with Western blotting at 2 weeks after injection of the recombinant systems (Fig. 2b and S3). AAV-IL4 mice were exposed to CMS for 4 weeks and the expression of IL4 in the hippocampus was confirmed again (Fig. 2c).

**Fig. 2.**
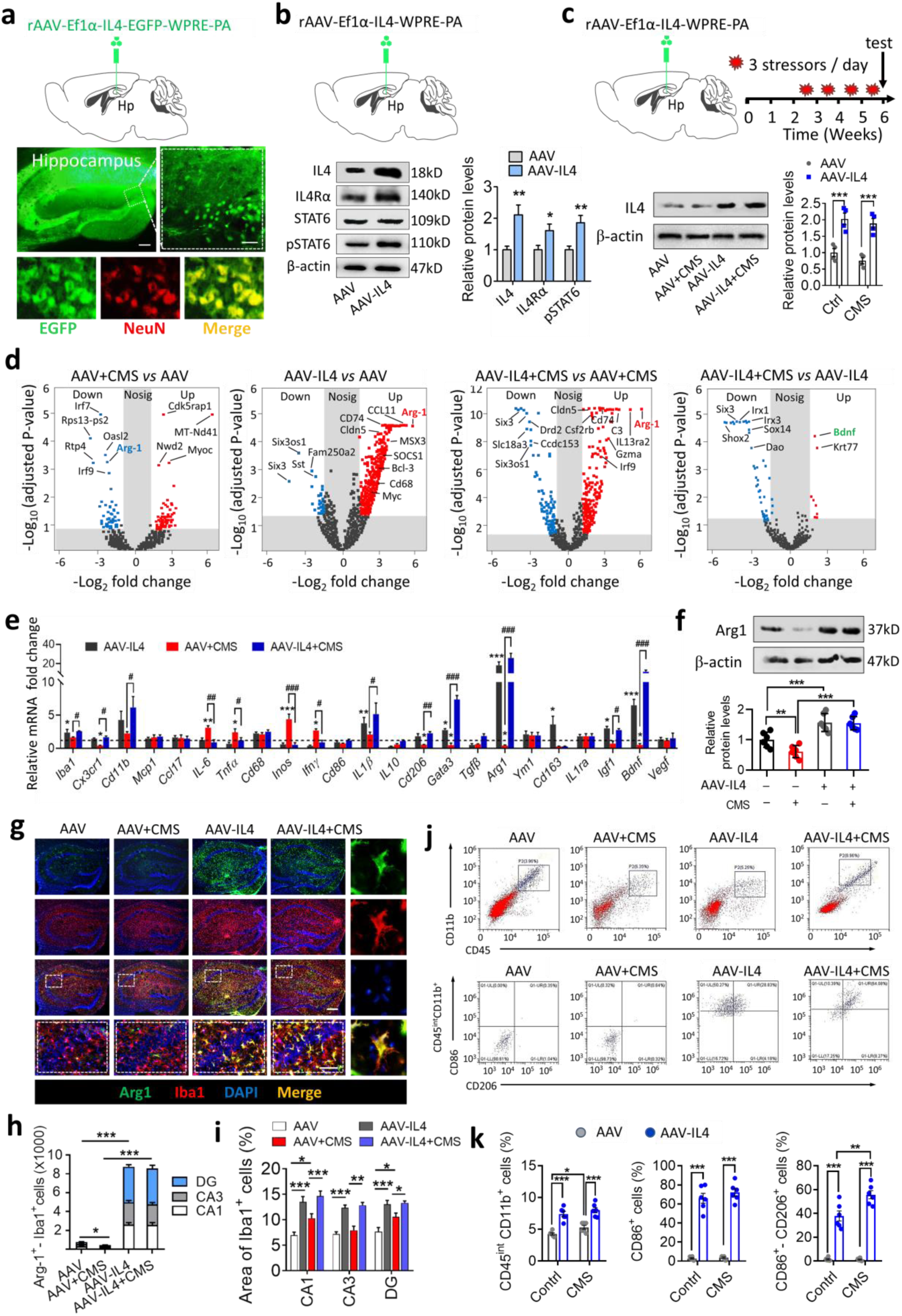
Overexpression of IL4 in the hippocampus enhanced Arg1^+^ microglia phenotypes and mediated neuroinflammatory responses to CMS paradigm. **a**, Representative fluorescence micrographs of hippocampus in mice injected with rAAV-Ef1α-IL4-EGFP-WPRE-PA. The lower illustration represents the co-localization of EGFP and NeuN. rAAV, Recombinant Adeno-Associated Virus; Ef1α, promoter driving target gene expression; IL4-EGFP, fusion of target gene (IL4) with enhanced green fluorescent protein; WPRE-pA, Woodchuck hepatitis virus post-transcriptional regulatory element. Scale bar, 100 μm (inset, 20 μm). **b**, Western blotting detection of IL4, IL4Rα, STAT6 and pSTAT6 in the hippocampus of mice after injecting AAV or AAV-IL4 for 2 weeks. The IL4 and IL4Rα are normalized to β-actin, the pSTAT6 is normalized to STAT6. Data are mean ± SEM (n = 6 per group). * P < 0.05, ** P < 0.01 (independent-samples *t*-tests). **c**, Western blotting detection of IL4 in the hippocampus of AAV or AAV-IL4 mice combined with CMS. The IL4 is normalized to β-actin. Data are mean ± SEM (n = 6 per group). *** P < 0.005 (two-way ANOVA with LSD test). **d**, Volcano maps indicate DEGs of AAV-IL4 or/and CMS-exposed brains. DEGs, differentially expressed genes. **e**, Quantitative PCR examination of the expression of several DEGs in the hippocampus. Data are mean ± SEM (n = 5–6 per group). * P < 0.05, ** P < 0.01, *** P < 0.005 (two-way ANOVA with LSD test). **f**, Western blotting revealed differences in the levels of Arg1 in the hippocampus of AAV-IL4 and/or CMS-exposed mice. Data are mean ± SEM (n = 6 per group). ** P < 0.01, *** P < 0.005 (two-way ANOVA with LSD test). **g**, Immunofluorescence micrographs show that Arg1 is localized to Iba1^+^ cells in the hippocampus of mice. Scale bar, 100 μm (inset, 20 μm). **h**, Quantification of Arg1^+^-Iba1^+^ cells in DG, CA1 and CA3 regions of the hippocampus. (n = 5–6 per group). * P < 0.05, *** P < 0.005 (two-way ANOVA with LSD test). **i**, Quantification of area of Iba1^+^ cells in DG, CA1 and CA3 of hippocampus. (n = 5– 6 per group). * P < 0.05, ** P < 0.01, *** P < 0.005 (two-way ANOVA with LSD test). **j**, Flow-cytometry of single cell suspension of the whole hippocampus for microglia (CD45^int^-CD11b^+^ cells), activated microglia (CD86^+^ cells) and anti-inflammatory microglia (CD86^+^-CD206^+^ cells). **k**, Quantification of area of CD45^int^-CD11b^+^ cells, CD86^+^ cells and CD86^+^-CD206^+^ cells. (n = 5–6 per group). * P < 0.05, ** P < 0.01, *** P < 0.005 (two-way ANOVA with LSD test).

Transcriptome sequencing was performed to explore the whole transcriptome levels of the hippocampus during AAV-IL4 and/or CMS treatment by RNA sequencing (RNA-seq) (Fig. S4). There were 65 genes differentially expressed in AAV + CMS mice when compared to AAV mice (31 showed upregulation and 34 downregulation). Most of these genes are involved in stress responses. There were 760 genes with differential expression in AAV-IL4 + CMS mice compared to AAV + CMS mice (597 showed upregulation and 163 downregulation). Most of these genes are involved in immunomodulation (Fig. 2d). Several of the genes were confirmed to be significantly and differentially expressed in the hippocampus by qPCR (Fig. 2e). Among them, Arg1 was the most significantly upregulated both at the gene and protein levels after AAV-IL4 injection with or without CMS treatment (Fig. 2f and Fig. S5). Further, the expression of Arg1 was strongly correlated with hippocampal IL4 levels (Fig. S5).

To determine the cell source of Arg1, we performed cell localization of Arg1. Immunohistochemical staining showed that Arg1 was located in hippocampal Iba1^+^ cells, and the number of Arg1^+^ microglia was significantly elevated in CA1, CA3 and DG of hippocampus in AAV-IL4 mice under both non-stress and stress conditions (Fig. 2g and 2h). Microglia in the hippocampus of AAV-IL4 mice showed obvious morphological changes, manifested by decreased processes and larger somata (Fig. 2i and S6). To assay the population of microglial subsets, flow-cytometry was performed on single cell suspensions of the whole hippocampus. The results revealed that microglia, in the form of CD45^int^-CD11b^+^ cells, increased significantly in CMS-exposed mice and AAV-IL4 mice, corroborating the presence of microglia proliferation. The CD86^+^-CD206^+^ microglia were significantly increased in the hippocampus of AAV-IL4 mice (Fig. 2j and 2k).

### Knockdown of microglial IL4Rα decreased Arg1^+^ microglia phenotype in the hippocampus

To confirm the effects of IL4 on microglial phenotype, knockdown of the microglial IL4 receptor was performed. The lentivirus encoding IL4Rα-targeting shRNA (LV-U6-LoxP-CMV-EGFP-LoxP-IL4Rα shRNA) was first synthesized. The transfection efficiency of LV-U6-LoxP-CMV-EGFP-LoxP-IL4Rα shRNA on microglia was then confirmed (Fig. S7). To test the efficiency of Cre recombinant enzyme, the hippocampi of CX_3_CR1^CreER^ mice were transfected with LV-U6-LoxP-CMV-EGFP-LoxP-IL4Rα shRNA, and then treated with tamoxifen to specifically recognize the LoxP site (Fig. S8a). After tamoxifen treatment for 1 week, the green fluorescent protein in microglia was cut efficiently (Fig. 3a). Next, we synthesized another lentivirus vector (LV-U6-LoxP-Stop-LoxP-IL4Rα shRNA) without the expression of EGFP to specifically downregulate microglial IL4Rα in hippocampus (Fig. S8b). After tamoxifen treatment for 1 week, the microglial IL4Rα, STAT6, pSTAT6 and Arg1 were significantly reduced in the hippocampus (Fig. 3b). We thus achieved IL4Rα shRNA specific targeting of hippocampal microglia (shIL4Rα/M).

**Fig. 3.**
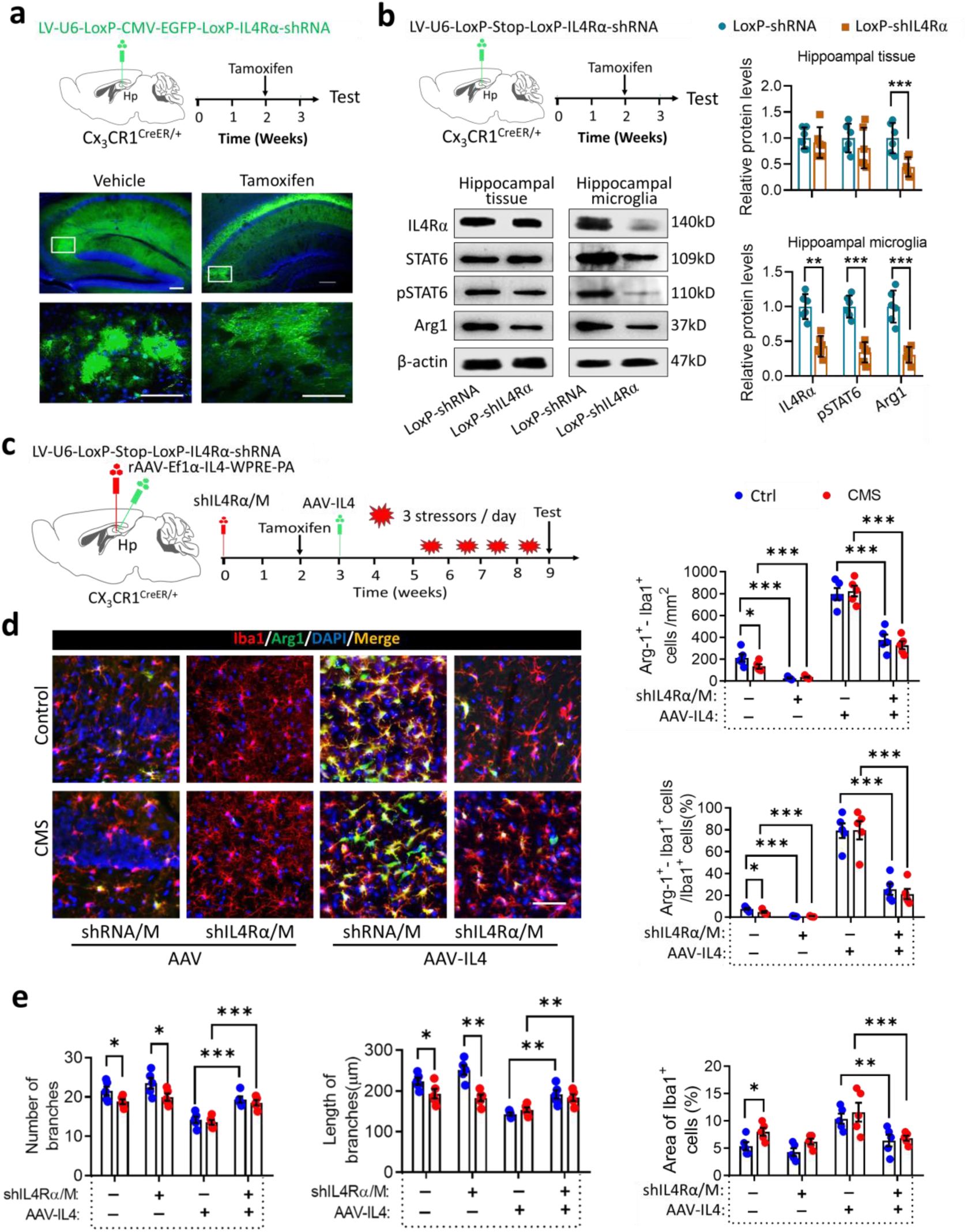
Knockdown of microglial IL4Rα decreased the Arg1^+^ microglia phenotype in the hippocampus. **a**, Efficiency test of Cre-recombinant enzyme in hippocampus of CX_3_CR1^CreER^ mice. Schematic and fluorescence micrographs of hippocampus in CX_3_CR1^CreER^ mice injected with LV-U6-LoxP-CMV-EGFP-LoxP-IL4Rα shRNA. The fluorescence micrographs show the expression of EGFP in microglia-like cells is suppressed by tamoxifen treatment. LV, lentivirus; U6, promoter driving target gene expression; CMV, promoter driving EGFP (green fluorescent protein); LoxP, specific recognition site of Cre-recombinant enzyme; IL4Rα shRNA, interference fragment for IL4Rα. Scale bar, 100 μm. **b,** Schematic of the strategy used to knockdown microglial IL4Rα and Western blotting of IL4Rα, STAT6, pSTAT6 and Arg1 in hippocampal tissue and microglia respectively. IL4Rα and Arg1 are normalized to β-actin, pSTAT6 is to normalized STAT6. Data are mean ± SEM (n = 6 per group). ** P < 0.01, *** P < 0.005 (independent-samples *t*-tests). **c**, Experimental timeline of the strategy used to knockdown microglial IL4Rα combined with overexpression of IL4 and CMS exposure. CX_3_CR1^CreER^ mice were injected stereotactically with LV-Loxp-shIL4Rα or LV-Loxp-shRNA into the hippocampus, allowed to recover for 2 weeks, then treated with tamoxifen to induce expression of microglial IL4Rα shRNA. Finally, the animals were injected with AAV-IL4, after 2 weeks, these mice were subjected to CMS consisting of exposure to three random stressors daily for 4 weeks. **d**, Effects of microglial IL4Rα knockdown in the hippocampus on the ratio of Arg1^+^ microglia. Scale bar, 20 μm. Data are mean ± SEM (n = 5 per group). * P < 0.05, *** P < 0.005 (two-way ANOVA with LSD test). **e**, Effects of knockdown of microglial IL4Rα before AAV-IL4 injection on the number, length of branches and area of Iba1^+^ cells in hippocampus of CMS-exposed CX_3_CR1^CreER^ mice. Data are mean ± SEM (n = 5-6 per group). * P < 0.05, ** P < 0.01, *** P < 0.005 (two-way ANOVA with LSD test).

Knockdown of microglial IL4Rα decreased the number of Arg1^+^ microglia phenotype in the hippocampus of AAV or AAV-IL4 *CX_3_CR1^CreER^* mice, under both non-stress and stress conditions (Fig. 3c and 3d). Remarkably, knockdown of microglial IL4Rα suppressed the proliferation of microglia, the decrease in length of branches and increase in area of Iba1^+^ cells in the hippocampus of AAV or AAV-IL4 *CX_3_CR1^CreER^* mice, under non-stress and stress conditions (Fig. 3e-3g).

### IL4-driven Arg1^+^ microglia enhanced hippocampal neurogenesis in CMS-exposed mice

GO enrichment analysis of differentially expressed genes in transcriptome sequencing showed that CMS affected expression of genes associated with stress response, apoptosis, and synaptic function. Overexpression of IL4 in the hippocampus of CMS-exposed mice resulted in changes related to neurogenesis, such as the nutrient metabolism, Notch and Wnt pathways (Fig. 4a).

**Fig. 4.**
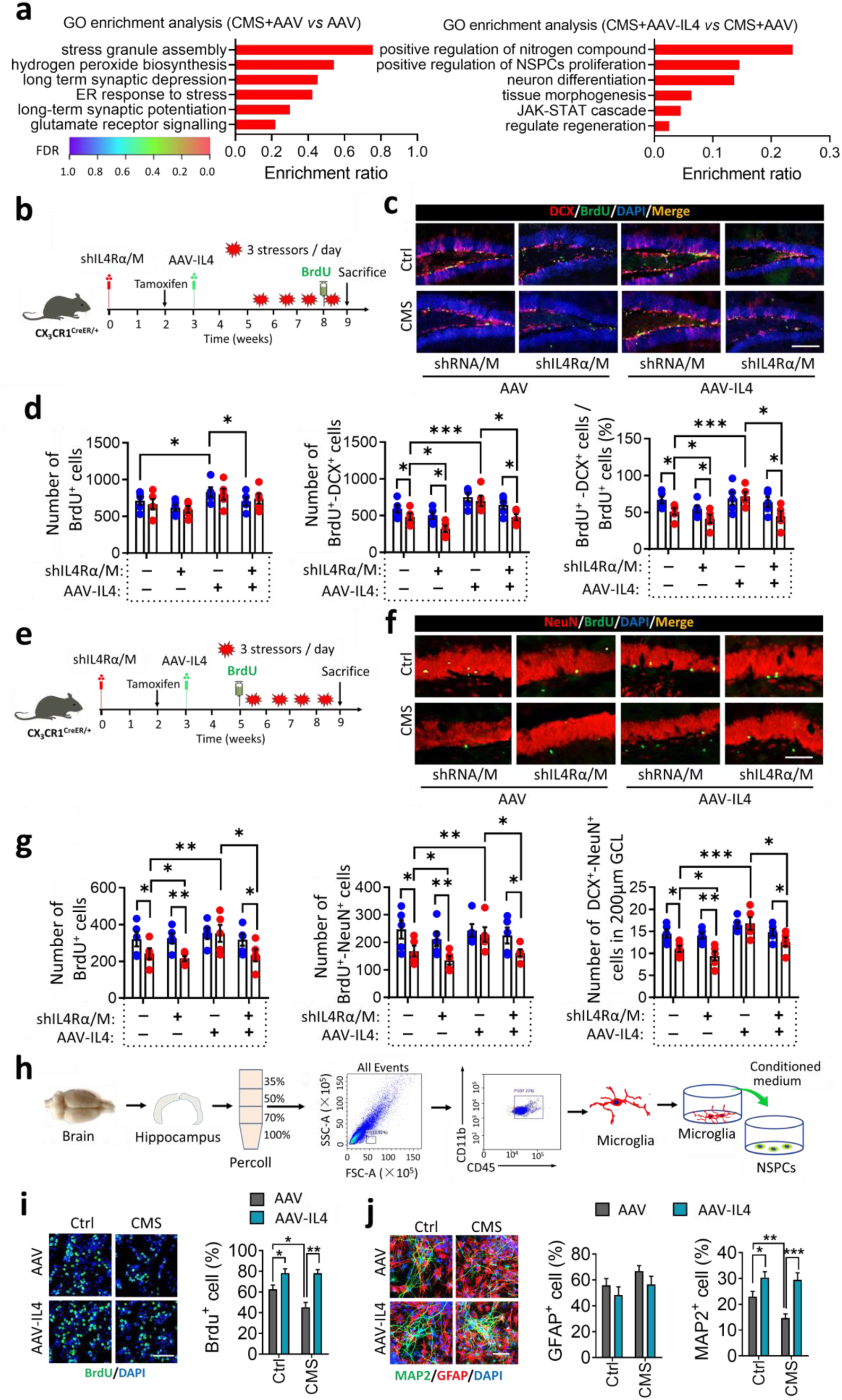
IL4-driven Arg1^+^ microglia enhanced hippocampal neurogenesis in CMS-exposed mice. **a**, GO enrichment analysis for DEGs. **b**, Experimental timeline of IL4 overexpression and/or microglial IL4Rα downregulation on NSPCs proliferation and differentiation in the hippocampus of CX_3_CR1^CreER^ mice. **c**, Representative fluorescence micrographs showing DCX expression and BrdU incorporation in the subgranular zone (SGZ). Scale bar, 100 μm. **d**, Quantification of total number of BrdU^+^ cells, total number of DCX^+^-BrdU^+^ cells and percentage of DCX^+^-BrdU^+^ cells out of all BrdU^+^ cells in the neurogenic zones. Data are mean ± SEM (n = 6 per group). * P < 0.05, ** P < 0.01, *** P < 0.005 (two-way ANOVA with LSD test). **e**, Experimental timeline of IL4 overexpression and/or microglial IL4Rα downregulation in the hippocampus (before IL4 overexpression) on NSPCs survival and maturation. **f**, Representative fluorescence micrographs illustrating NeuN expression and BrdU incorporation in the SGZ. Scale bar, 100 μm. **g**, Quantification of total number of BrdU^+^ cells, BrdU^+^-NeuN^+^ cells and DCX^+^-NeuN^+^ cells in DG. Data are mean ± SEM (n = 5 per group). * P < 0.05, ** P < 0.01, *** P < 0.005 (two-way ANOVA with LSD test). **h**, Schematic describing NSPCs culture in conditioned medium from microglia isolated from hippocampus of AAV, AAV + CMS, AAV-IL4 and AAV-IL4 + CMS mice. **i**, Effects of conditioned medium from microglia isolated from hippocampus on NSPCs proliferation. Proliferation was monitored by quantifying the percentage of BrdU^+^ cells. Scale bar, 50 μm. Data are mean ± SEM (n = 6 per group). * P < 0.05, ** P < 0.01 (two-way ANOVA with LSD test). **j**, Representative micrographs of NSPCs cultured for 7 days in conditioned culture medium from microglia isolated from hippocampus. Cells were immunolabeled with GFAP to identify astrocytes, and MAP2 to identify neurons. Percentages of GFAP^+^ cells and MAP2^+^ cells are quantified. Scale bar, 30 μm. Data are mean ± SEM (n = 6 per group). * P < 0.05, ** P < 0.01, *** P < 0.005 (two-way ANOVA with LSD test).

We examined hippocampal neurogenesis, including proliferation, differentiation, survival and maturity of NSPCs. The results showed that CMS-exposed mice displayed a striking reduction in neuronal differentiation, as shown by a lower number of BrdU^+^-DCX^+^ cells, and a slower neuronal differentiation rate compared with unstressed controls. Overexpression of IL4 with AAV-expression system in the hippocampus reversed CMS-induced deficits in neurogenesis. Meanwhile, knockdown of microglial IL4Rα decreased BrdU^+^-DCX^+^ cells and the percentage of BrdU^+^-DCX^+^ cells out of total BrdU^+^ cells in AAV or AAV-IL4 mice (Fig. 4b – 4d and Fig. S9).

To assay the extent of survival and maturity of proliferative cells, the proliferating cells were labeled with BrdU before CMS exposure (Fig. 4e). Reduced numbers of surviving cells (BrdU^+^), mature neurons (BrdU^+^-NeuN^+^) and maturing neurons (DCX^+^-NeuN^+^) were found in DG of CMS-exposed mice. Overexpression of IL4 reversed the CMS-induced decrease in the numbers of BrdU^+^ cells, BrdU^+^-NeuN^+^ cells and DCX^+^-NeuN^+^ cells in DG. Knockdown of microglial IL4Rα decreased BrdU^+^ cells, BrdU^+^-NeuN^+^ cells and DCX^+^-NeuN^+^ cells in AAV or AAV-IL4 mice (Fig. 4f, 4g and Fig. S9).

Considering that microglia can modulate neurogenesis, we collected the conditioned medium of microglia isolated from the hippocampus of AAV mice or AAV-IL4 mice under stress or non-stress conditions to culture NSPCs (Fig. 4h). In NSPCs proliferation assays, the microglial conditioned medium (M-CM) from AAV mice with CMS exposure decreased neurosphere size and percentage of BrdU^+^ cells when compared the M-CM from AAV mice without CMS exposure. The opposite effects were observed with M-CM from AAV-IL4 mice without CMS. The M-CM from AAV-IL4 mice with CMS exposure increased neurosphere size and the percentage of BrdU^+^ cells when compared the M-CM from mice injected with AAV and with CMS exposure (Fig. 4i and Fig. S10). This suggested that CMS-induced microglia suppressed the proliferation of NSPCs, and IL4-driven microglia promoted the proliferation of NSPCs in the hippocampus of CMS-exposed mice.

In NSPCs differentiation assays, we found that M-CM from mice injected with AAV and with CMS exposure decreased the percentage of MAP2^+^ cells and NG2^+^ cells when compared to the M-CM from AAV mice without CMS exposure. This result suggested that CMS-induced microglia suppressed the differentiation of NSPCs into neuron or oligodendrocyte. The M-CM from AAV-IL4 mice with or without CMS exposure increased the percentage of MAP2^+^ cells and BrdU^+^-MAP2^+^ cells (Fig. 4j and Fig. S10). These data suggest that IL4-driven microglia promote differentiation of NSPCs to neuron and survival of immature neurons. The M-CM from IL4-stimulated primary microglia had a similar effect on proliferation and differentiation of NSPCs. However, there were no significant effects on proliferation and differentiation of NSPCs after they were treated with IL4 when compared with phosphate-buffered saline (PBS) (Fig. S11).

### IL4-driven microglia enhance neurogenesis via a BDNF-dependent pathway

Next we explored how IL4-driven Arg-1^+^ microglia may promote neurogenesis. In the transcriptome study, the expression of *Bdnf* was significantly upregulated in the brain of AAV-IL4 + CMS mice. Concurrently, nutrient component metabolism and BDNF receptor signaling-related genes were enriched to a high degree (Fig. 5a). This suggested that IL4-driven microglia may promote hippocampal neurogenesis through BDNF signaling pathways. We confirmed that BDNF concentration was maintained at a high level in the hippocampus of AAV-IL4 mice in the presence or absence of CMS treatment (Fig. 5b). The signal of BDNF receptor (TrkB) in the hippocampus also was strengthened by overexpression of IL4 (Fig. S12). Immunohistochemistry showed that BDNF accumulated in large quantities around Iba1^+^ cells in the hippocampus of AAV-IL4 mice (Fig. 5c). Notably, we found BDNF widely distributed in both intracellular and extracellular compartments of Arg1^+^ microglia (Fig. 5d). These results suggested that IL4-driven microglia could synthesize and secrete more BDNF. Knockdown of microglial IL4Rα significantly reduced BDNF expression in the hippocampus of AAV-IL4 mice (Fig. 5e and 5f). Correlation analysis showed that there was a significant positive correlation between the number of BrdU^+^-DCX^+^ cells and the concentration of BDNF in the hippocampus (Fig. 5g).

**Fig. 5.**
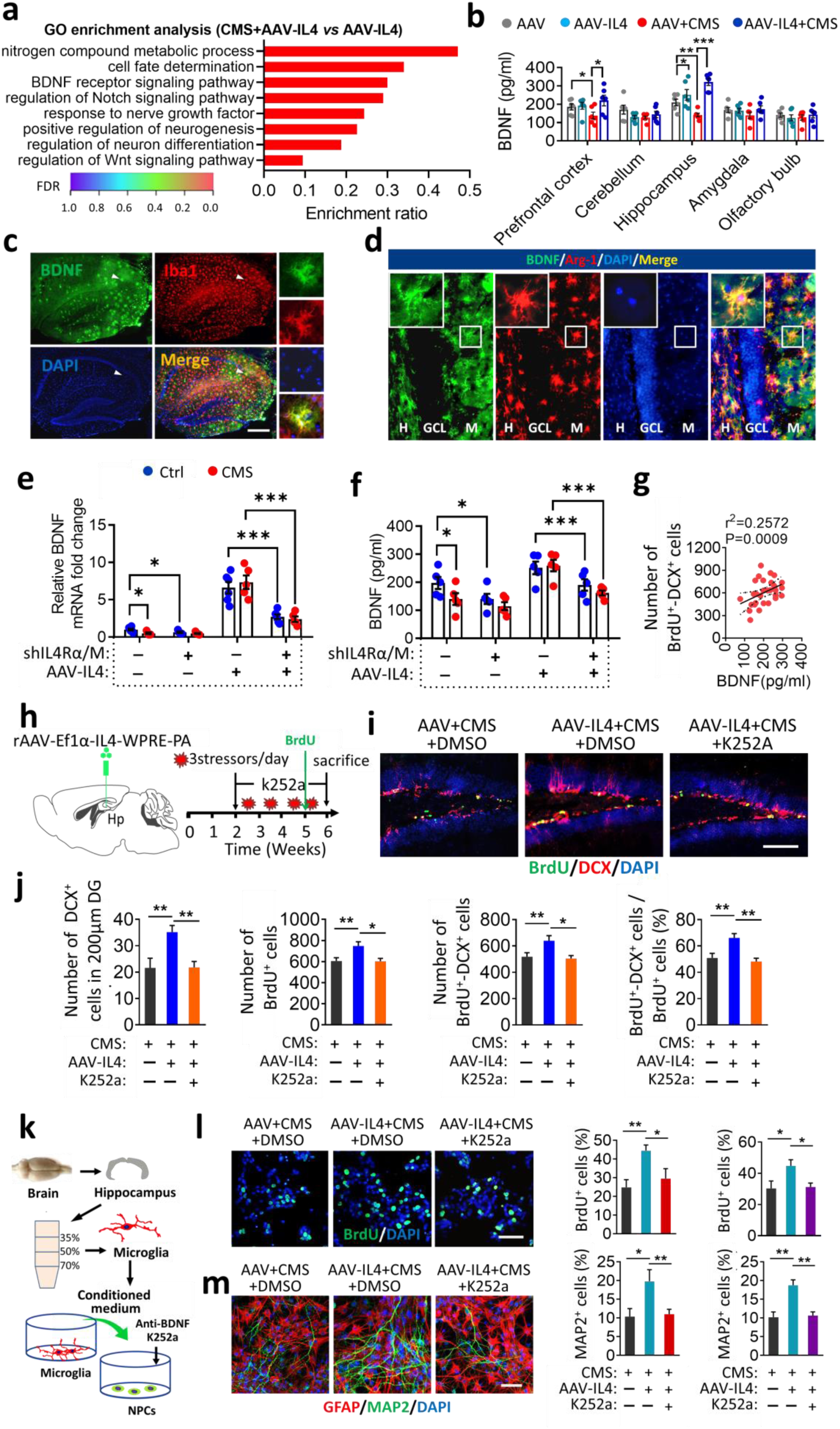
IL4-driven microglia modulate adult hippocampal neurogenesis in a BDNF-dependent manner. **a**, GO enrichment analysis for DEGs. **b**, ELISA to assay BDNF levels in prefrontal cortex, cerebellum, hippocampus, amygdala and olfactory bulb. Data are mean ± SEM (n = 5–7 per group). * P < 0.05, ** P < 0.01, *** P < 0.005 (two-way ANOVA with LSD test). **c**, Double immunohistochemical staining of BDNF and Iba1 in the hippocampus of AAV-IL4 mice. Scale bar, 50 μm. **d**, Double immunohistochemical staining of BDNF and Arg1 in the hippocampus of AAV-IL4 mice. Scale bar, 20 μm. **e** and **f**, Effects of CMS, knockdown of microglial IL4Rα and overexpression of IL4 on BDNF mRNA and protein levels in the hippocampus. Data are mean ± SEM (n = 5–6 per group). * P < 0.05, ** P < 0.01, *** P < 0.005 (two-way ANOVA with LSD test). **g**, Correlation between total number of DCX^+^-BrdU^+^ cells per DG and percentage of hippocampal BDNF level. Each circle represents one mouse (n = 6 per group). **h**, Schematic representation of k252a treatment to block the BDNF/TrkB pathway in AAV-IL4 mice. **i**, Representative fluorescence micrographs showing DCX expression and BrdU incorporation in the SGZ. Scale bar, 100 μm. **j**, Quantification of the number of DCX^+^ cells, total number of BrdU^+^ cells and DCX^+^-BrdU^+^ cells and percentage of DCX^+^-BrdU^+^ cells out of all BrdU^+^ cells in the neurogenic zones. Data are mean ± SEM (n = 5–6 per group). * P < 0.05, ** P < 0.01 (two-way ANOVA with LSD test). **k**, Schematic of treatment with anti-BDNF antibody (anti-BDNF) or BDNF receptor antagonist (K252a) to block the BDNF/TrkB pathway in NSPCs cultured in microglial conditioned medium. **l**, Effects of anti-BDNF antibody or K252a on NSPCs proliferation (BrdU^+^ cells) in the presence the conditioned culture medium from microglia isolated from hippocampus of AAV-IL4 + CMS mice. Scale bar, 20 μm. Data are mean ± SEM (n = 6 per group, 5 replicates per sample). * P < 0.05, ** P < 0.01 (one-way ANOVA with LSD test). **m**, Effects of anti-BDNF antibody or K252a on NSPCs differentiation in the presence of conditioned culture medium from microglia isolated from hippocampus of AAV-IL4 + CMS mice. Scale bar, 10 μm (I) and 20 μm (J). Data are mean ± SEM (n = 6 per group, 5 replicates per sample). * P < 0.05, ** P < 0.01 (one-way ANOVA with LSD test).

To determine if BDNF was necessary and sufficient for IL4-rescued neurogenesis, AAV-IL4 mice were treated with K252a to block the TrkB pahway (Fig. 5h). We found that K252a treatment blocked the increase in DCX^+^ cells, BrdU^+^ cells and BrdU^+^-DCX^+^ cells in the hippocampus of AAV-IL4 mice subjected to the CMS paradigm (Fig. 5i and 5j). BDNF/TrkB signaling was blocked *in vitro* with either an anti-BDNF neutralizing antibody or the BDNF receptor antagonist K252a (Fig. 5k). As expected, both reagents were able to block the increase of NSPCs proliferation, differentiation, and migration when NSPCs were cultured in conditioned medium from microglia isolated from hippocampus of AAV-IL4 mice exposed to CMS (Fig. 5l, 5m and Fig S13). These results suggest that BDNF is necessary for the pro-neurogenesis effects of IL4-driven microglia in CMS-exposed mice.

### IL4-driven Arg1^+^ microglia protected mice from stress-induced depressive-like behaviors

We examined the effects of IL4 overexpression on resistance of mice to stress-induced depression using a series of behavioral tests. After 4 weeks of CMS-exposure, AAV mice showed less sucrose preference, longer immobility time in TST and FST, longer latency to feed in NSFT, and shorter time in the center in OFT than non-stressed control animals. As expected, these depressive-like behaviors were significantly absent in AAV-IL4 mice exposed to CMS (Fig. 6a and Fig. S14). Through correlation analysis, we found that sucrose preference and immobility time in FST were positively and negatively correlated, respectively, with the percentage of Arg1^+^ microglia, BDNF concentration and BrdU^+^-DCX^+^ cells in the hippocampus (Fig. 6b). Remarkably, after knockdown of microglial IL4Rα in the hippocampus, AAV-IL4 CX_3_CR1^CreER^ mice showed depressive-like behaviors after CMS exposure, and AAV CX_3_CR1^CreER^ mice showed higher stress vulnerability than control mice to the CMS paradigm (Fig. 6c, Fig S15 and Fig. S16). As mentioned above, IL4-driven microglia modulate hippocampal neurogenesis in a BDNF-dependent manner in CMS-treated mice. Our experiments showed that either K252a or TMZ treatment abolished the antidepressant effects of IL4-driven microglia based on the SPT, FST, TST, and NSF (Fig. 6d, 6e and Fig. S17). These results suggest that IL4-driven Arg1^+^ microglia enhance hippocampal neurogenesis and stress resistance of CMS-exposed mice via a BDNF-dependent pathway.

**Fig. 6.**
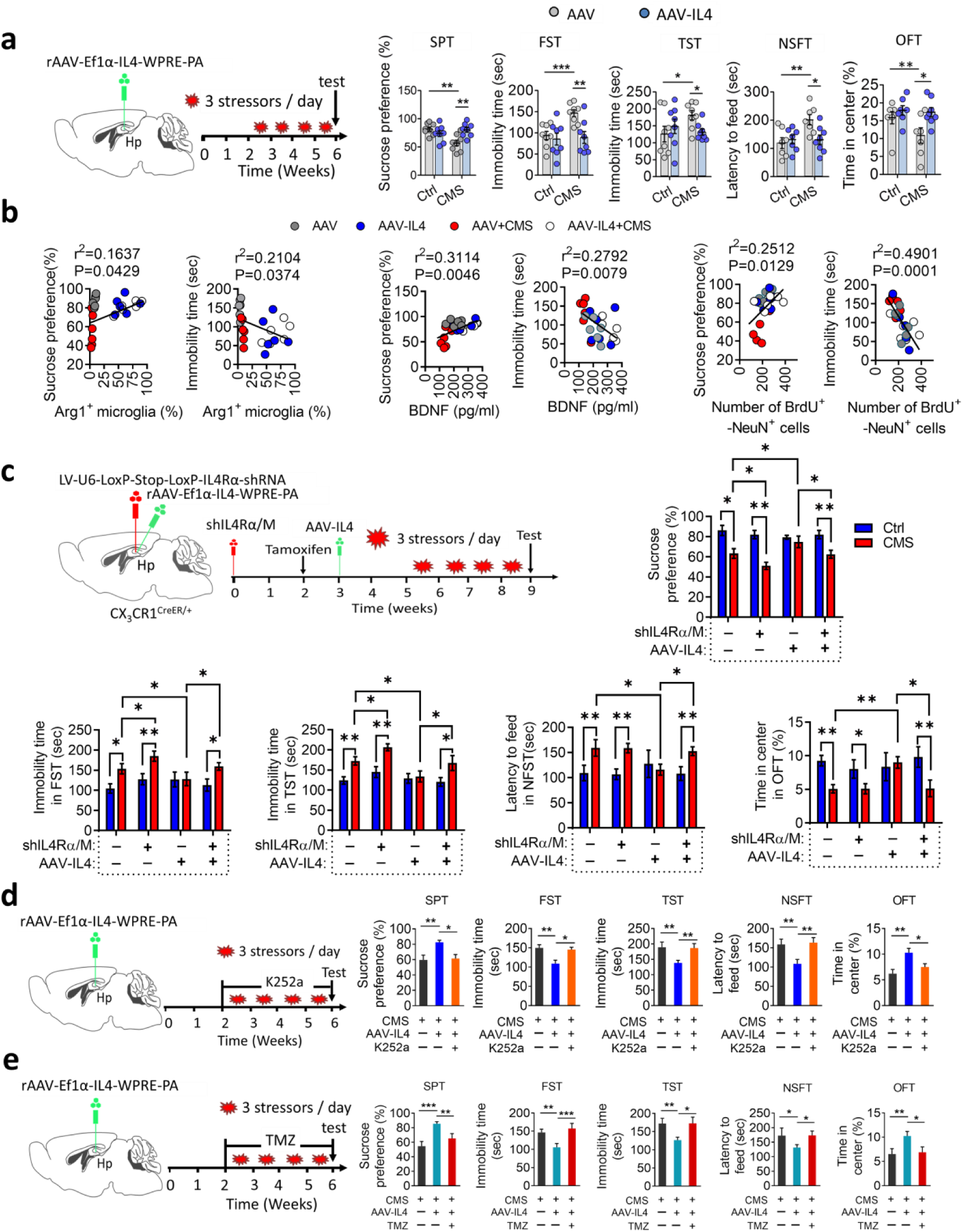
IL4-driven Arg1^+^ microglia protected mice from stress-induced depressive-like behaviors. **a**, Effects of overexpression of IL4 in the hippocampus on stress-induced depressive-like behaviors. Mice were injected stereotactically with rAAV-IL4 or rAAV, allowed to recover for 2 weeks, then subjected to a CMS protocol consisting of exposure to three random stressors daily for 4 weeks. The mice were assessed by SPT, FST, TST, NSFT and OFT. Ctrl, non-stress mice; CMS, CMS-exposed mice; AAV, AAV injection; AAV-I-4, AAV-IL4 injection. SPT, sucrose preference test; FST, forced swim test; TST, tail suspension test; NSFT, novelty-suppressed feeding test; OFT, open field test. Data are mean ± SEM (n = 8–10 per group). * P < 0.05, ** P < 0.01, *** P < 0.005 (two-way ANOVA with LSD test). **b,** Correlations between sucrose preference and percentage of Arg1^+^ microglia in hippocampus, immobility time in FST and percentage of Arg1^+^ microglia in hippocampus, sucrose preference and concentration of BDNF in hippocampus, immobility time in FST and concentration of BDNF in hippocampus, sucrose preference and number of BrdU^+^-DCX^+^ cells in DG, immobility time in FST and number of BrdU^+^-DCX^+^ cells in DG. Each circle represents one mouse (n = 5 per group). **c**, Effects of microglial IL4Rα downregulation (before IL4 overexpression) on stress-induced depressive-like behaviors, as assessed by SPT, TST, FST, NSFT and OFT). Data are mean ± SEM (n = 8-10 per group). * P < 0.05, ** P < 0.01 (two-way ANOVA with LSD test). **d**, Assessment of stress-induced depressive-like behaviors in AAV-IL4 mice when treated with k252a to block the BDNF/K252a pathway. Data are mean ± SEM (n = 8– 10 per group). * P < 0.05, ** P < 0.01 (one-way ANOVA with LSD test). **e**, Assessment of stress-induced depressive-like behaviors in AAV-IL4 mice when treated with TMZ (temozolomide) to block neurogenesis. Data are mean ± SEM (n = 8–10 per group). * P < 0.05, ** P < 0.01, *** P < 0.005 (one-way ANOVA with LSD test).

## Discussion

Microglia are endowed with phenotypic plasticity, which can be stimulated by different cytokines to regulate physiological responses and behavioral outcomes during stress ^34, 35^. In particular, IL4 induces Arg1^+^ microglial phenotype and promotes neural repair ^19^. The present study provides, to the best of our knowledge, the first direct evidence that IL4-driven microglia are essential for hippocampal neurogenesis and that they can mediate the resilience to chronic stress. The promotion of neurogenesis by IL4-driven Arg1^+^ microglia was associated with upregulation of BDNF in the hippocampus. Thus, our study demonstrates that IL4-driven Arg1^+^ microglia ameliorate chronic stress-induced depressive-like behaviors in mice through BDNF-dependent neurogenesis.

The underlying immune environment alters the interaction between stress and inflammatory processes, thus contributing to the risk of depressive symptoms ^36, 37^. In our study we found that the levels of IL4 were decreased in the hippocampus of HS mice, but increased in LS mice. Vulnerability to chronic stress is associated with dysregulation of pro- and anti-inflammatory cytokines, in particular an ineffective IL4 pathway ^21^. The reduction of IL4 levels can render individuals more vulnerable to stress and more likely to engage in depressive-like behaviors. IL4 deficiency increases stress vulnerability and risk of mood disorders ^20^. Our results, together with evidence from the literature ^23, 38, 39^, suggest that levels of hippocampal IL4 vary inversely with stress vulnerability.

Upregulation of IL4 in the hippocampus protected mice from stress-induced depressive-like behaviors, consistent with the finding that IL4 overexpression has antidepressant effects ^21^. Our transcriptome analysis showed that CMS leads to decreased expression of genes related to immune regulation (Arg1, Irf7, Irf9, etc.) and increased expression of genes related to stress response (Myoc, Cdk5rap1, etc.). These results suggest that stress-induced depression-like behaviors are associated with neuroimmune regulation ^10^. Overexpression of IL4 in the hippocampus of CMS-exposed mice leads to increased expression of a large number of immune-related genes, suggesting that increased immune regulation in the hippocampus can increase stress resistance. Among them, Arg1 is the most significantly upregulated gene, and it is co-localized with microglia. Hence, overexpression of IL4 could significantly increase the number of Arg1^+^ microglia in hippocampus.

IL4-induced Arg1^+^ microglia can be thought of a special phenotype of beneficial microglia ^40, 41^. These cells regulate inflammatory processes in the brain and enhance neurite growth, thereby exhibiting neuroprotective effects ^27, 42, 43^. In our study, CMS-exposed mice with a lower level of hippocampal IL4 showed fewer Arg1^+^ microglia and higher levels of proinflammatory mediators such as IL1β, TNFα, IFNγ, and iNOS. In contrast, overexpression of IL4 in the hippocampus significantly increased Arg1^+^ microglia and suppressed levels of proinflammatory mediators. These findings suggest that IL4-driven Arg1^+^ microglia reduce inflammation in the hippocampus of CMS-exposed mice by reducing the synthesis and secretion of pro-inflammatory mediators. Further, the results from flow-cytometry analysis of single cell suspensions from the whole hippocampus showed that microglia, in the form of CD45^int^-CD11b^+^ cells, increased significantly in AAV-IL4 mice. These data indicate that IL4-induced Arg1^+^ microglia are characterized by high proliferative activity. Furthermore, CD86^+^-CD206^+^ microglia were significantly increased in the hippocampus of AAV-IL4 mice, suggesting that IL4 mediates neuroprotective effects in the CNS mainly by driving the phenotypic shift of microglia. Significantly, knockdown of microglial IL4Rα decreased the number of Arg1^+^ microglia and abrogated the antidepressant effects of IL4 overexpression, suggesting that IL4-driven Arg1^+^ microglia play a key role in protecting from CMS-induced depression-like behaviors.

Microglia constitute a prominent cell population within the hippocampal neurogenic niche ^44^. Our results confirmed that CMS resulted in pro-inflammatory activation of microglia and inhibition of hippocampal neurogenesis. The transcriptome sequencing showed that overexpression of IL4 in CMS-exposed mice induced microglial pro-neurogenic phenotype and mitigated stress-induced neurogenesis impairment. Knockdown of microglial IL4Rα of mice decreased the number of Arg1^+^ microglia, blocked IL4-rescued hippocampal neurogenesis and stress resistance in CMS-treated mice. The data from *in vitro* assays showed that microglia isolated from hippocampus of CMS-exposed mice suppressed NSPCs proliferation and neurogenesis. Meanwhile, microglia isolated from hippocampus of AAV-IL4 mice were sufficient to enhance NSPCs proliferation and neuronal differentiation. These effects were not observed when IL4 was added directly to NSPCs, which indicated that IL4 rescued the impairment of adult neurogenesis in CMS-exposed mice in a microglia-dependent manner ^45 24, 46, 47^. Inhibition of neurogenesis *in vivo* with TMZ abolished the antidepressant effects of IL4-activated microglia in CMS-treated mice. Adult hippocampal neurogenesis appears to occur in humans as well as in rodents, although this idea is still controversial ^48–50^. The relatively small number of new neurons in adult brains of humans and rodents is strongly linked to the pathogenesis and remission of neuropsychiatric disorders ^51^. These findings suggest that microglia-mediated pro-neurogenesis underlies resistance to stress-induced depression.

IL4 drove microglia to adopt a phenotype of alternative activation in CMS-exposed mice, associated with higher levels of BDNF release. Although BDNF/TrkB signaling was reduced in the hippocampus of CMS-exposed mice, the transcriptome sequencing with GO enrichment analysis showed BDNF signaling-related genes were enriched to a high degree in AAV-IL4 mice. Furthermore, we found that abundant BDNF is produced by Arg1^+^ microglia. Knockdown of microglial IL4Rα decreased the BDNF levels in the hippocampus of AAV-IL4 mice. Blocking of BDNF/TrkB signaling with k252a blocked IL4-rescued hippocampal neurogenesis and abolished stress resistance in AAV-IL4 mice. In addition, blocking BDNF/TrkB signaling *in vitro* also eliminated microglia-mediated neurogenesis. These findings suggest that the pro-neurogenesis effects of IL4-driven Arg1^+^ microglia depend on BDNF/TrkB signaling. Because microglia are myeloid in origin, different from nerve cells, it was believed that microglia influence neuronal behavior through non-contact-dependent pathways ^52^. BDNF is an important regulator of hippocampal neurogenesis and is necessary for neurobehavioral plasticity ^53, 54^. Our results suggest that the pro-neurogenesis effects and therefore the resiliency to stress of IL4-driven Arg1^+^ microglia depend on BDNF/TrkB signaling.

## Conclusions

The present study identified critical roles of Arg1^+^ microglia-mediated pro-neurogenesis in the modulation of vulnerability or resistance of mice to stress. Reprogramming microglia with IL4 can rescue hippocampal neurogenesis in stressed mice by increasing BDNF signaling. These findings indicate that regulation of microglial function is a potential therapeutic strategy for the treatment of mood disorders.

## Acknowledgments

This work was supported by the National Natural Science Foundation of China (81571174, 81701308, U1808204) and the 863 project (2015AA020505). We thank Ruotian Jiang in Department of Anesthesiology, West China Hospital, Sichuan University, and Hedong Li in Department of Biology, Pennsylvania State University, for useful discussion and valuable suggestions. We are grateful to Prof. Zujun Yang for providing materials and assistance with immunohistochemistry, and to A. Chapin Rodríguez for help in revising the manuscript.

## References

1. Hanson, N.D., Owens, M.J. & Nemeroff, C.B. Depression, antidepressants, and neurogenesis: a critical reappraisal. Neuropsychopharmacology : official publication of the American College of Neuropsychopharmacology 36, 2589–2602 (2011).

2. Anacker, C., et al. Hippocampal neurogenesis confers stress resilience by inhibiting the ventral dentate gyrus. Nature 559, 98–102 (2018).

3. Teh, D.B., Ishizuka, T. & Yawo, H. Regulation of later neurogenic stages of adult-derived neural stem/progenitor cells by L-type Ca2+ channels. Development, growth & differentiation 56, 583–594 (2014).

4. Eisch, A.J. & Petrik, D. Depression and hippocampal neurogenesis: a road to remission? *Science (New York*, N.Y.) 338, 72–75 (2012).

5. Wang, H., et al. Norbin ablation results in defective adult hippocampal neurogenesis and depressive-like behavior in mice. Proceedings of the National Academy of Sciences of the United States of America 112, 9745–9750 (2015).

6. Rotheneichner, P., et al. Hippocampal neurogenesis and antidepressive therapy: shocking relations. 2014, 723915 (2014).

7. Yirmiya, R., Rimmerman, N. & Reshef, R. Depression as a microglial disease. Trends in neurosciences 38, 637–658 (2015).

8. Shigemoto-Mogami, Y., Hoshikawa, K., Goldman, J.E., Sekino, Y. & Sato, K. Microglia enhance neurogenesis and oligodendrogenesis in the early postnatal subventricular zone. The Journal of neuroscience : the official journal of the Society for Neuroscience 34, 2231–2243 (2014).

9. Kreisel, T., et al. Dynamic microglial alterations underlie stress-induced depressive-like behavior and suppressed neurogenesis. Molecular psychiatry 19, 699–709 (2014).

10. Zhang, J., et al. Salvianolic acid B promotes microglial M2-polarization and rescues neurogenesis in stress-exposed mice. Brain, behavior, and immunity 66, 111–124 (2017).

11. Hill, A.S., Sahay, A. & Hen, R. Increasing Adult Hippocampal Neurogenesis is Sufficient to Reduce Anxiety and Depression-Like Behaviors. Neuropsychopharmacology : official publication of the American College of Neuropsychopharmacology 40, 2368–2378 (2015).

12. Ziv, Y., et al. Immune cells contribute to the maintenance of neurogenesis and spatial learning abilities in adulthood. Nature neuroscience 9, 268–275 (2006).

13. Littlefield, A.M., Setti, S.E., Priester, C. & Kohman, R.A. Voluntary exercise attenuates LPS-induced reductions in neurogenesis and increases microglia expression of a proneurogenic phenotype in aged mice. Journal of neuroinflammation 12, 138 (2015).

14. Parkhurst, C.N., et al. Microglia promote learning-dependent synapse formation through brain-derived neurotrophic factor. Cell 155, 1596–1609 (2013).

15. Coull, J.A., et al. BDNF from microglia causes the shift in neuronal anion gradient underlying neuropathic pain. Nature 438, 1017–1021 (2005).

16. Reshef, R., Kreisel, T., Beroukhim Kay, D. & Yirmiya, R. Microglia and their CX3CR1 signaling are involved in hippocampal- but not olfactory bulb-related memory and neurogenesis. Brain, behavior, and immunity 41, 239–250 (2014).

17. Pluchino, S. & Peruzzotti-Jametti, L. Interleukin-4 induced 1 (IL4I1) promotes central nervous system remyelination. Brain : a journal of neurology 139, 3052–3054 (2016).

18. Gadani, S.P., Cronk, J.C., Norris, G.T. & Kipnis, J. IL-4 in the brain: a cytokine to remember. Journal of immunology (Baltimore, Md. : 1950) 189, 4213–4219 (2012).

19. Zhao, X., et al. Neuronal Interleukin-4 as a Modulator of Microglial Pathways and Ischemic Brain Damage. 35, 11281–11291 (2015).

20. Uceyler, N., Topuzoglu, T., Schiesser, P., Hahnenkamp, S. & Sommer, C. IL-4 deficiency is associated with mechanical hypersensitivity in mice. PloS one 6, e28205 (2011).

21. Wachholz, S., et al. Interleukin-4 is a participant in the regulation of depressive-like behavior. Behavioural brain research 326, 165–172 (2017).

22. Park, H.J., et al. IL-4 Inhibits IL-1beta-Induced Depressive-Like Behavior and Central Neurotransmitter Alterations. Mediators of inflammation 2015, 941413 (2015).

23. Han, A., et al. IL-4/10 prevents stress vulnerability following imipramine discontinuation. Journal of neuroinflammation 12, 197 (2015).

24. Butovsky, O., et al. Microglia activated by IL-4 or IFN-gamma differentially induce neurogenesis and oligodendrogenesis from adult stem/progenitor cells. Molecular and cellular neurosciences 31, 149–160 (2006).

25. Yu, Z. & Sun, D. MSX3 Switches Microglia Polarization and Protects from Inflammation-Induced Demyelination. 35, 6350–6365 (2015).

26. Pepe, G., Calderazzi, G., De Maglie, M., Villa, A.M. & Vegeto, E. Heterogeneous induction of microglia M2a phenotype by central administration of interleukin-4. Journal of neuroinflammation 11, 211 (2014).

27. Cherry, J.D., Olschowka, J.A. & O’Banion, M.K. Arginase 1+ microglia reduce Abeta plaque deposition during IL-1beta-dependent neuroinflammation. Journal of neuroinflammation 12, 203 (2015).

28. Nasca, C., Bigio, B., Zelli, D., Nicoletti, F. & McEwen, B.S. Mind the gap: glucocorticoids modulate hippocampal glutamate tone underlying individual differences in stress susceptibility. Molecular psychiatry 20, 755–763 (2015).

29. Nam, B.Y., et al. Double transduction of a Cre/LoxP lentiviral vector: a simple method to generate kidney cell-specific knockdown mice. American journal of physiology. Renal physiology 309, F1060–1069 (2015).

30. Cao, X., et al. Astrocyte-derived ATP modulates depressive-like behaviors. Nature medicine 19, 773–777 (2013).

31. Norden, D.M., Fenn, A.M., Dugan, A. & Godbout, J.P. TGFbeta produced by IL-10 redirected astrocytes attenuates microglial activation. Glia 62, 881–895 (2014).

32. Fenn, A.M., Henry, C.J., Huang, Y., Dugan, A. & Godbout, J.P. Lipopolysaccharide-induced interleukin (IL)-4 receptor-alpha expression and corresponding sensitivity to the M2 promoting effects of IL-4 are impaired in microglia of aged mice. Brain, behavior, and immunity 26, 766–777 (2012).

33. Rubio-Perez, J.M. & Morillas-Ruiz, J.M. A review: inflammatory process in Alzheimer’s disease, role of cytokines. TheScientificWorldJournal 2012, 756357 (2012).

34. Sato, K. Effects of Microglia on Neurogenesis. Glia 63, 1394–1405 (2015).

35. Fenn, A.M., et al. Immune activation promotes depression 1 month after diffuse brain injury: a role for primed microglia. Biological psychiatry 76, 575–584 (2014).

36. Ehrlich, K.B., Miller, G.E., Rohleder, N. & Adam, E.K. Trajectories of relationship stress and inflammatory processes in adolescence. Development and psychopathology 28, 127–138 (2016).

37. Pearson-Leary, J., et al. Inflammation and vascular remodeling in the ventral hippocampus contributes to vulnerability to stress. Translational psychiatry 7, e1160 (2017).

38. Menard, C., Pfau, M.L., Hodes, G.E. & Russo, S.J. Immune and Neuroendocrine Mechanisms of Stress Vulnerability and Resilience. Neuropsychopharmacology : official publication of the American College of Neuropsychopharmacology 42, 62–80 (2017).

39. Lee, H.J., Park, H.J., Starkweather, A., An, K. & Shim, I. Decreased Interleukin-4 Release from the Neurons of the Locus Coeruleus in Response to Immobilization Stress. 2016, 3501905 (2016).

40. Ma’arif, B., Agil, M. & Laswati, H. The enhancement of Arg1 and activated ERbeta expression in microglia HMC3 by induction of 96% ethanol extract of Marsilea crenata Presl. leaves. Journal of basic and clinical physiology and pharmacology (2020).

41. Cai, W., et al. STAT6/Arg1 promotes microglia/macrophage efferocytosis and inflammation resolution in stroke mice. JCI insight 4 (2019).

42. Toedebusch, C.M., et al. Arginase-1 expressing microglia in close proximity to motor neurons were increased early in disease progression in canine degenerative myelopathy, a model of amyotrophic lateral sclerosis. Molecular and cellular neurosciences 88, 148–157 (2018).

43. Zhao, Q., et al. Phenotypic dysregulation of microglial activation in young offspring rats with maternal sleep deprivation-induced cognitive impairment. Scientific reports 5, 9513 (2015).

44. Diaz-Aparicio, I., et al. Microglia actively remodel adult hippocampal neurogenesis through the phagocytosis secretome. The Journal of neuroscience : the official journal of the Society for Neuroscience (2020).

45. You, Z., et al. Pro- and anti-inflammatory cytokines expression in rat’s brain and spleen exposed to chronic mild stress: involvement in depression. Behavioural brain research 225, 135–141 (2011).

46. Paintlia, A.S., Paintlia, M.K., Singh, I. & Singh, A.K. IL-4-induced peroxisome proliferator-activated receptor gamma activation inhibits NF-kappaB trans activation in central nervous system (CNS) glial cells and protects oligodendrocyte progenitors under neuroinflammatory disease conditions: implication for CNS-demyelinating diseases. Journal of immunology (Baltimore, Md. : 1950) 176, 4385–4398 (2006).

47. Bhattarai, P., et al. IL4/STAT6 Signaling Activates Neural Stem Cell Proliferation and Neurogenesis upon Amyloid-beta42 Aggregation in Adult Zebrafish Brain. Cell reports 17, 941–948 (2016).

48. Sorrells, S.F., et al. Human hippocampal neurogenesis drops sharply in children to undetectable levels in adults. Nature 555, 377–381 (2018).

49. Boldrini, M., et al. Human Hippocampal Neurogenesis Persists throughout Aging. Cell stem cell 22, 589–599.e585 (2018).

50. Kempermann, G., et al. Human Adult Neurogenesis: Evidence and Remaining Questions. Cell stem cell 23, 25–30 (2018).

51. Yun, S., Reynolds, R.P., Masiulis, I. & Eisch, A.J. Re-evaluating the link between neuropsychiatric disorders and dysregulated adult neurogenesis. Nature medicine 22, 1239–1247 (2016).

52. Eid, R.S., et al. Early and late effects of maternal experience on hippocampal neurogenesis, microglia, and the circulating cytokine milieu. Neurobiology of aging 78, 1–17 (2019).

53. Liu, P.Z. & Nusslock, R. Exercise-Mediated Neurogenesis in the Hippocampus via BDNF. Frontiers in neuroscience 12, 52 (2018).

54. Leal, G., Bramham, C.R. & Duarte, C.B. BDNF and Hippocampal Synaptic Plasticity. Vitamins and hormones 104, 153–195 (2017).

