## Supplementary Tables for "IL4-driven microglia modulate stress resilience through BDNF-dependent neurogenesis"

**Table S1. Details of q-PCR primers**

| Gene | Primer sequences |
|---|---|
| *β-actin* | Forward: 5’-CCGTGAAAAGATGACCCAGATC-3’<br>Reverse: 5’-CACAGCCTGGATGGCTACGT-3’ |
| *Cd11b* | Forward: 5’-CACAATGGATGGCTTGATGGA-3’<br>Reverse: 5’-CGTCCACGCAGTCCGGTAAAA-3’ |
| *Gfap* | Forward: 5’-TGCTGGAGGGCGAAGAAA-3’<br>Reverse: 5’-CGGATCTGGAGGTTGGAGAA-3’ |
| *Cd68* | Forward: 5’-CCACAGGCAGCACAGTGGACA-3’<br>Reverse: 5’-TCCACAGCAGAAGCTTTGGCCC-3’ |
| *Cd86* | Forward: 5’-ACGATGGACCCCAGATGCACCA-3’<br>Reverse: 5’-GCGTCTCCACGGAAACAGCA-3’ |
| *Infγ* | Forward: 5’-CGGCACAGTCATTGAAAGCCTA-3’<br>Reverse: 5’-GTTGCTGATGGCCTGATTGTC-3’ |
| *IL1β* | Forward: 5’-CCAGCAGGTTATCATCATCATCC-3’<br>Reverse: 5’-CTCGCAGCAGCACATCAAC-3’ |
| *IL2* | Forward:5’-CCTGAGCAGGATGGAGAATTACA-3’<br>Reverse: 5’-TCCAGAACATGCCGCAGAG-3’ |
| *IL6* | Forward: 5-ACCGCTATGAAGTTCCTCTC-3’<br>Reverse: 5’-CTCTGTGAAGTCTCCTCTCC-3’ |
| *IL12* | Forward: 5’-GCCCTCTCTCTCCTCTTGCT-3’<br>Reverse: 5’-GTCTGCCTCTTTTGGTCAGG-3’ |
| *Mcp1* | Forward: 5’-TTAACGCCCCCACTCACCTGCTG-3’<br>Reverse: 5’-GCTTCTTTGGGACACCTGCTGC-3’ |
| *Gata3* | Forward:5’-GCCATTCGTACATGGAAGC-3’<br>Reverse: 5’-CGGAGGGTAAACGGACAGAG-3’ |
| *Tnfα* | Forward:5’-TACTGAACTTCGGGGTGATTGGTCC-3’<br>Reverse: 5’-CAGCCTTGTCCCTTGAAGAGAACC-3’ |
| *Inos* | Forward: 5’-ACAACAGGAACCTACCAGCTCA-3’<br>Reverse: 5’-GATGTTGTAGCGCTGTGTGTCA-3’ |
| *Ccl17* | Forward: 5’-CTCAGTTCATCCACGGCATAC-3’<br>Reverse: 5’-GACAAGGCTCACCATCATCG-3’ |
| *Cd163* | Forward: 5’-CGGCACAGTCATTGAAAGCCTA-3’<br>Reverse: 5’-GTTGCTGATGGCCTGATTGTC-3’ |
| *IL4* | Forward: 5’-TCTCGAATGTACCAGGAGCCATATC-3’<br>Reverse: 5’-AGCACCTTGGAAGCCCTACAGA-3’ |
| *IL10* | Forward: 5’-TGGCCCAGAAATCAAGGAGC-3’<br>Reverse: 5’-CAGCAGACTCAATACACACT-3’ |
| *Tgfβ* | Forward: 5’-GACCGCAACAACGCCATCTA-3’<br>Reverse: 5’-GGCGTATCAGTGGGGGTCAG-3’ |
| *Fizz1* | Forward: 5’-TCCCAGTGAATACTGATGAGA-3’<br>Reverse: 5’-CCACTCTGGATCTCCCAAGA-3’ |

| | |
|---|---|
| *Arg1* | Forward: 5'-AGACAGCAGAGGAGGTGAAGAG-3'<br>Reverse: 5'-CGAAGCAAGCCAAGGTTAAAGC-3' |
| *Mmr* | Forward: 5'-AGTTGGGTTCTCCTGTAGCCCAA-3'<br>Reverse: 5'-ACTACTACCTGAGCCCACACCTGCT-3' |
| *Ym1* | Forward: 5'-CATTCAGTCAGTTATCAGATTCC-3'<br>Reverse: 5'-AGTGAGTAGCAGCCTTGG-3' |
| *IL1ra* | Forward: 5'-CCAGCTCATTGCTGGGTACT-3'<br>Reverse: 5'-TTCTCAGAGCGGATGAAGGT-3' |
| *IL13* | Forward: 5'-CAATTGCAATGCCATCTACAGGAC-3'<br>Reverse: 5'-CGAAACAGTTGCTTTGTGTAGCTGA-3' |
| *C3* | Forward: 5'-TCGGATGACAAGGTCACCCT-3'<br>Reverse: 5'-AAGTGGCTCTCGTACCACAG-3' |
| *Cx3cr1* | Forward: 5'-GCGCCGCAGCAAGTCT-3'<br>Reverse: 5'-TTGAGAGCTGACTGCTGGAAGTT-3' |
| *Igf1* | Forward: 5'-GGTGGTTTATGAATGGTT-3'<br>Reverse: 5'-AGGGTGTGTCTAATGGAG-3' |
| *Bdnf* | Forward: 5'-GAGCTGAGCGTGTGTGACAG-3'<br>Reverse: 5'-CGCCAGCCAATTCTCTTTTTGC-3' |
| *Vegf* | Forward: 5'-CCATGAACTTTCTGCTCTTC-3'<br>Reverse: 5'-GGTGAGAGGTCTAGTTCCCGA-3' |
| *Cxcl10* | Forward: 5'-CATTCAGTCAGTTATCAGATTCC-3'<br>Reverse: 5'-AGTGAGTAGCAGCCTTGG-3' |
| *Iba1* | Forward: 5'-CGAGCTCATTGGTGGGTACT-3'<br>Reverse: 5'-TTCTCAGAGAGGATGCAGGT-3' |

**Table S2. Details of Anti-body and dilution rate.**

| Anti-Body | Tedia Company Inc | Dilution Rate |
|---|---|---|
| Iba1 (Goat anti-mouse) | Abcam | 1:400 |
| GFAP (Rat anti-mouse) | Cell Signaling Technology | 1:500 |
| iNOS (Rabbit anti-mouse) | Abcam | 1:50 |
| Arg-1 (Rabbit anti-mouse) | Abcam | 1:200 |
| CD68 (Rabbit anti-mouse) | Abcam | 1:300 |
| DCX (Goat anti-mouse) | Santa Cruz | 1:400 |
| NeuN (Rabbit anti-mouse) | Cell Signaling Technology | 1:800 |
| MAP2 (Rabbit anti-mouse) | Cell Signaling Technology | 1:500 |
| BrdU (Rat anti-mouse) | Cell Signaling Technology | 1:400 |
| NG2 (Rabbit anti-mouse) | Cell Signaling Technology | 1:500 |
| ßIII-tubulin (Rat anti-mouse) | Cell Signaling Technology | 1:200 |
| BDNF(Rat anti-mouse) | Abcam | 1:300 |
