## Supplementary Figures for "IL4-driven microglia modulate stress resilience through BDNF-dependent neurogenesis"

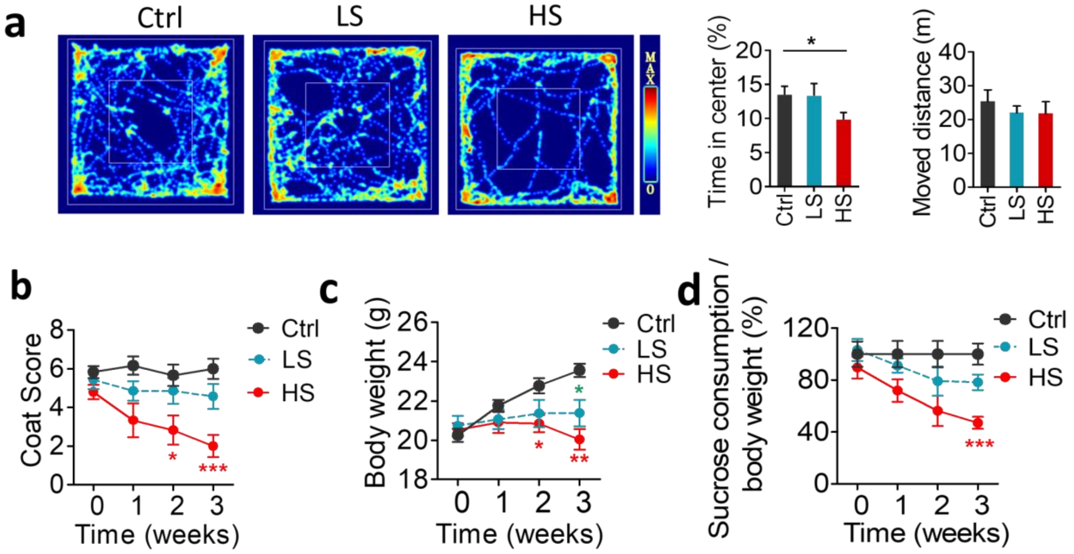


**Fig. S1. Validation of LS and HS mice**

**a**, Open field test for control, HS and LS mice. Quantification of the percentage of time spent in the center, and the travel distance of control, HS and LS mice in the OFT.

**b,** Weekly coat scores of control, HS and LS mice.

**c**, Weekly body weight of control, HS and LS mice.

**d**, Weekly sucrose consumption of control, HS and LS mice. Sucrose consumption was normalized to body weight. Data are mean ± SEM (n = 11–20 per group). * $P < 0.05$, ** $P < 0.01$, *** $P < 0.005$ (one-way ANOVA with LSD test).

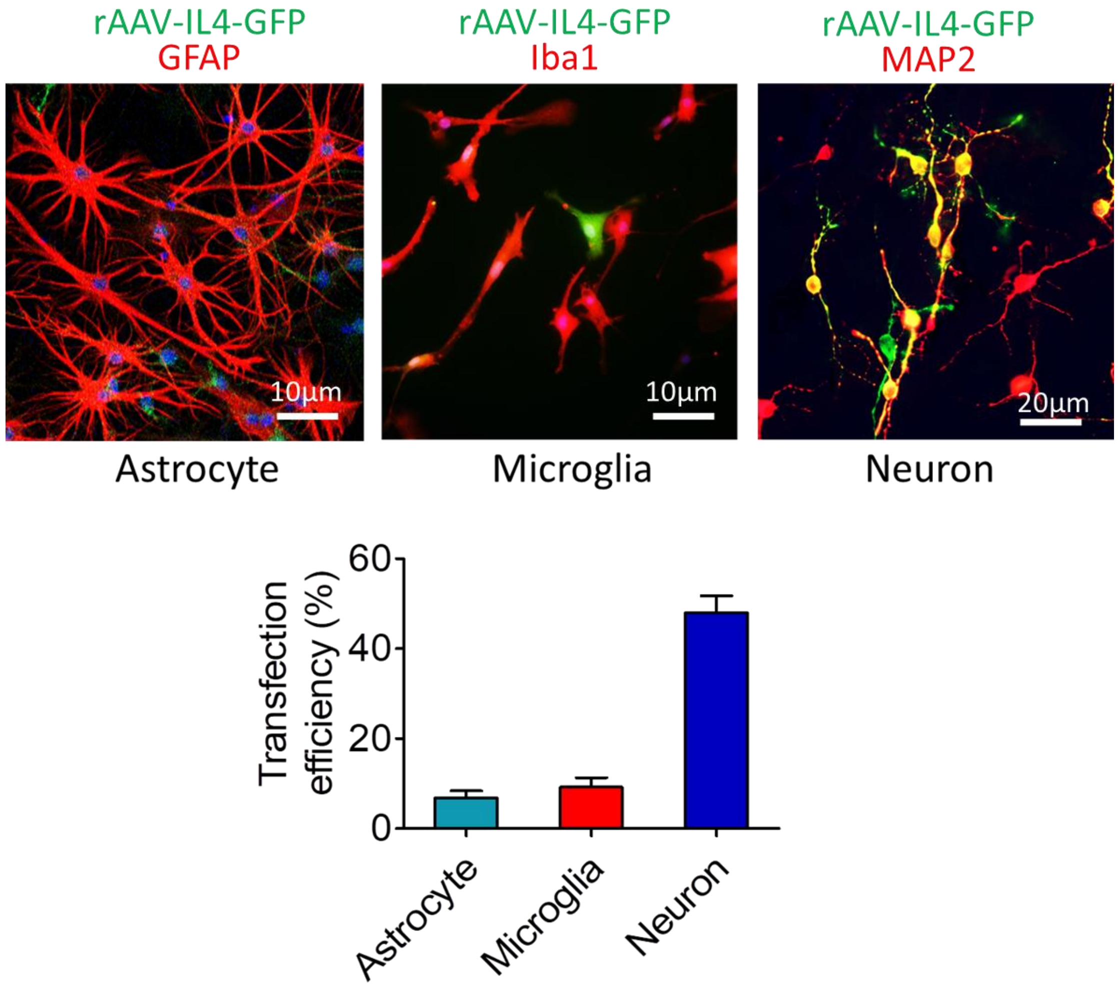


**Fig. S2. Evaluation of the transfection efficiency of AAV-IL4-EGFP in astrocytes, microglia and neurons *in vitro***

Cultured astrocytes, microglia and neurons were incubated with AAV-IL4-EGFP for 48 h, and the transfection efficiency was evaluated by computing the percentage of GFAP$^+$-EGFP$^+$ cells, Iba1$^+$-EGFP$^+$ cells and MAP2$^+$-EGFP$^+$ cells (n = 4).

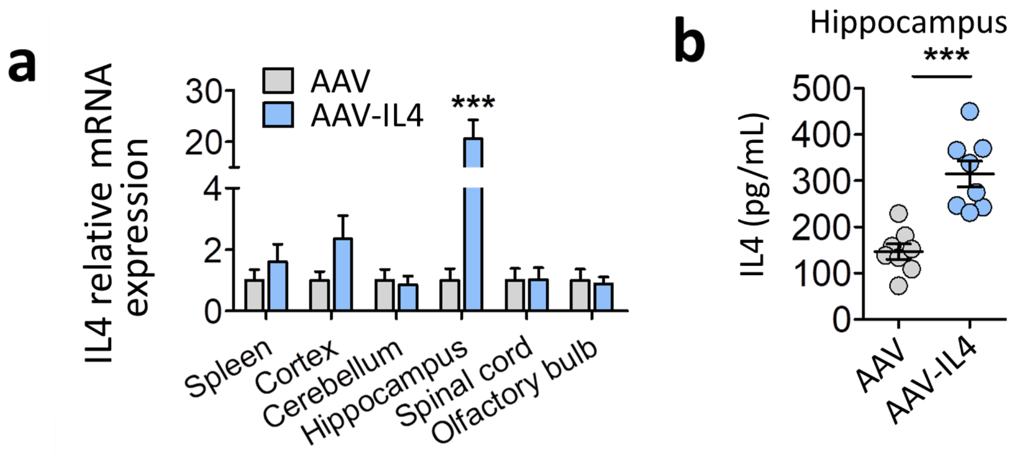


**Fig. S3. AAV-IL4 injection upregulated the expression of IL4 in the hippocampus**

**a**, mRNA expression of IL4 in cortex, cerebellum, amygdala, hippocampus and olfactory bulb of AAV mice and AAV-IL4 mice. Data are mean ± SEM (n = 4–5 per group).

**b**, ELISA examination of IL4 protein concentration in the hippocampus of AAV mice and AAV-IL4 mice. Data are mean ± SEM (n = 8 and 5 per group, respectively). * $P < 0.05$, ** $P < 0.01$, *** $P < 0.005$ (unpaired *t* test).

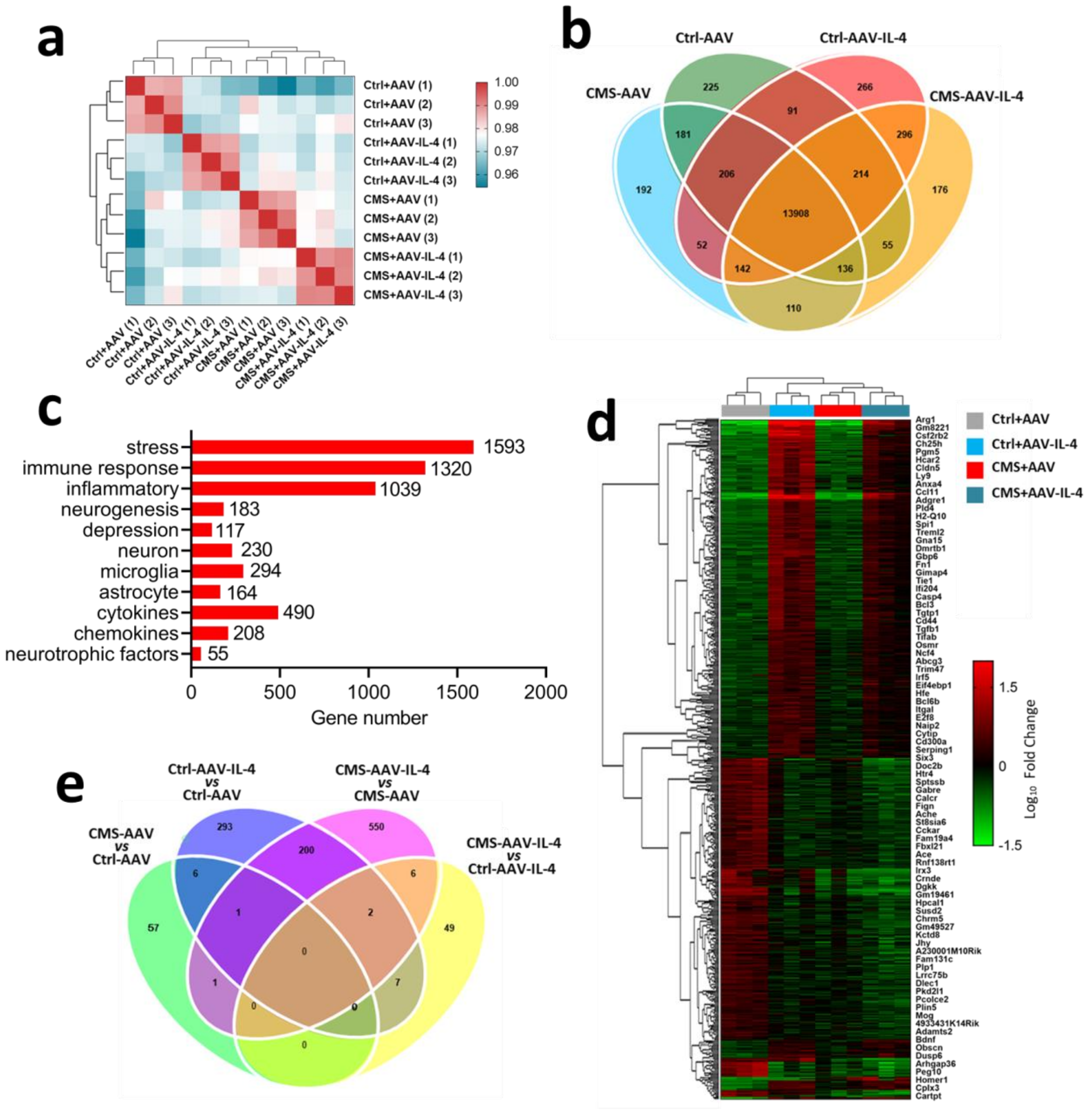


**Fig. S4. Transcriptome data analysis**

**a**, Sample correlation cluster analysis.

**b,** Wayne chart of differentially expressed genes in different groups.

**c**, Enrichment analysis of differentially expressed genes related to stress response.

**d**, Stratified clustering analysis of differentially expressed genes.

**e**, Wayne chart of differentially expressed genes in the comparison groups.

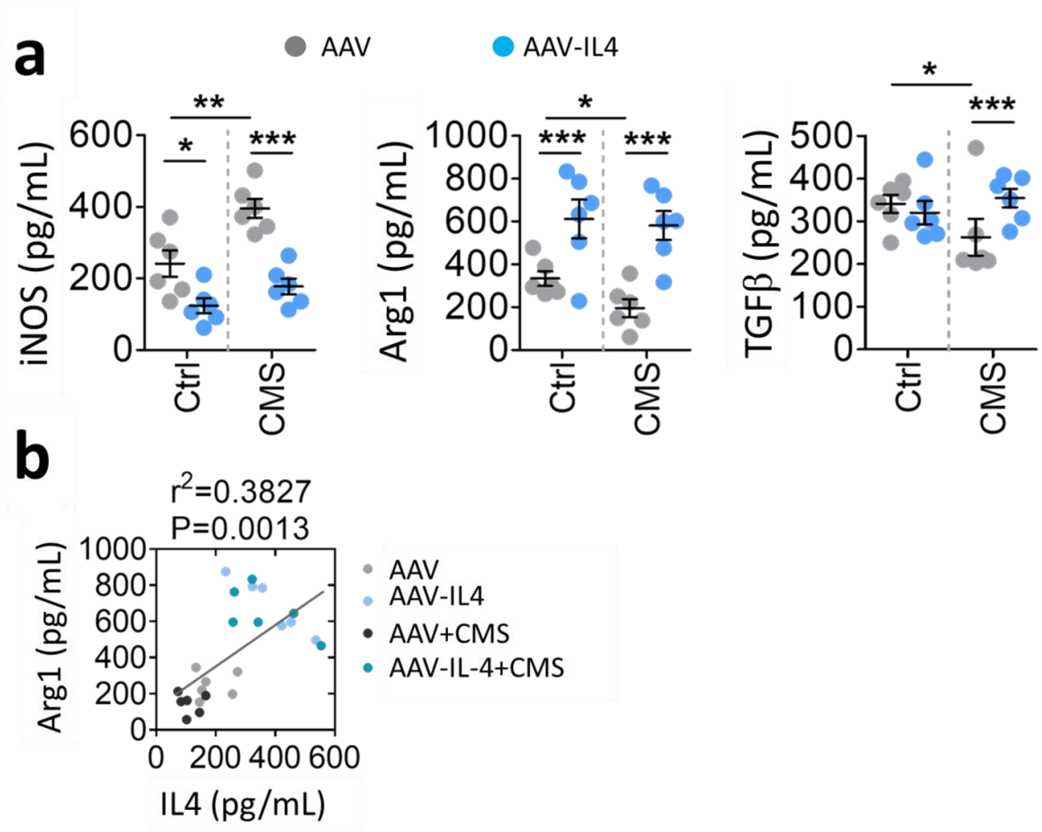


**Fig. S5. Differentially expressed cytokines were analyzed for protein levels**

**a**, ELISA examination of the protein concentrations of iNOS, Arg1 and TGFβ in the hippocampus of AAV mice, AAV-IL4 mice, AAV + CMS mice and AAV-IL4 + CMS mice. Data are mean ± SEM (n = 5 per group). * $P < 0.05$, ** $P < 0.01$, *** $P < 0.005$ (two-way ANOVA with LSD test).

**b,** Correlation between concentration of Arg1 and concentration of IL4 in the hippocampus of AAV mice, AAV-IL4 mice, AAV + CMS mice and AAV-IL4 + CMS mice.

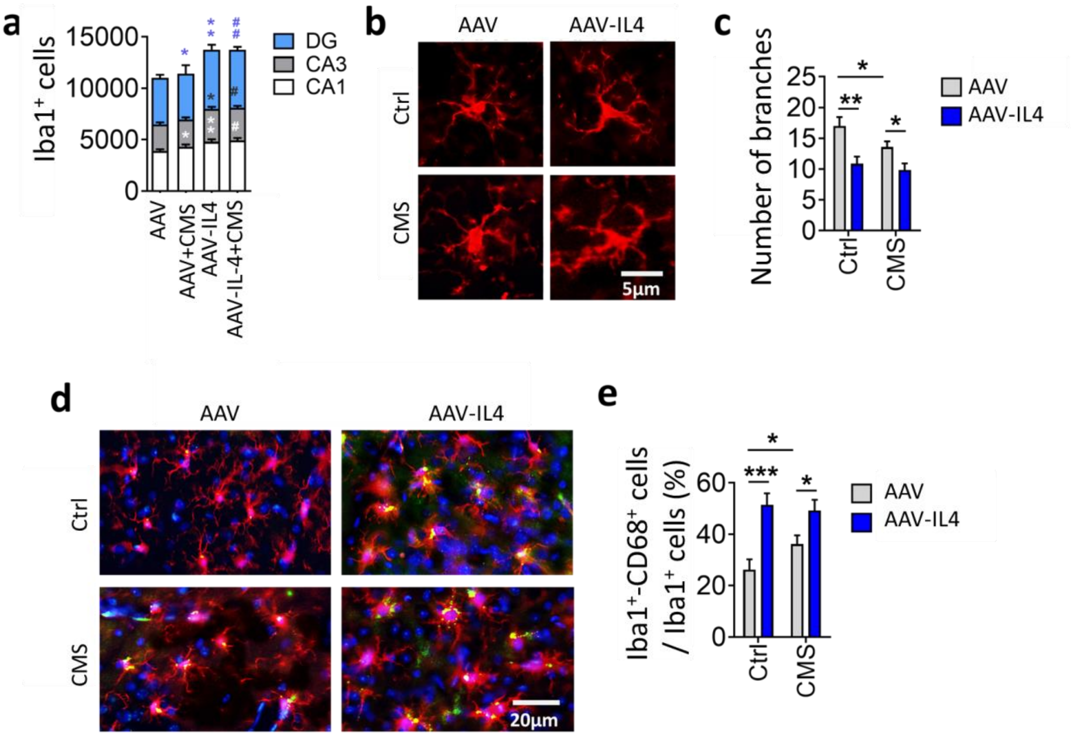


**Fig. S6. Effects of overexpression of IL4 on microglial activation in the hippocampus**

**a**, Quantification of the number of Iba1$^+$ cells in CA1, CA3 and DG of AAV, AAV-CMS, AAV-IL4 and AAV-IL4-CMS mice. Data are mean ± SEM (n = 5–6 per group).

**b**, Representative fluorescence micrographs illustrating microglial morphology in the hippocampus of AAV, AAV + CMS, AAV-IL4 and AAV-IL4 + CMS mice. Scale bar, 5 μm.

**c**, Quantification of the number of branches per Iba1$^+$ cell in the hippocampus of AAV, AAV + CMS, AAV-IL4 and AAV-IL4 + CMS mice. Data are mean ± SEM (n = 5–6 per group).

**d**, Representative fluorescence micrographs of microglia in the hippocampus of AAV, AAV + CMS, AAV-IL4 and AAV-IL4 + CMS mice. Activated microglia were stained immunohistochemically for Iba1 and CD68. Scale bar, 20 μm.

**e**, Quantification of the percentage of Iba1$^+$-CD68$^+$ cells in the hippocampus of AAV, AAV + CMS, AAV-IL4 and AAV-IL4 + CMS mice. Data are mean ± SEM (n = 5–6 per group). * P < 0.05, *** P < 0.005 (two-way ANOVA with LSD test).

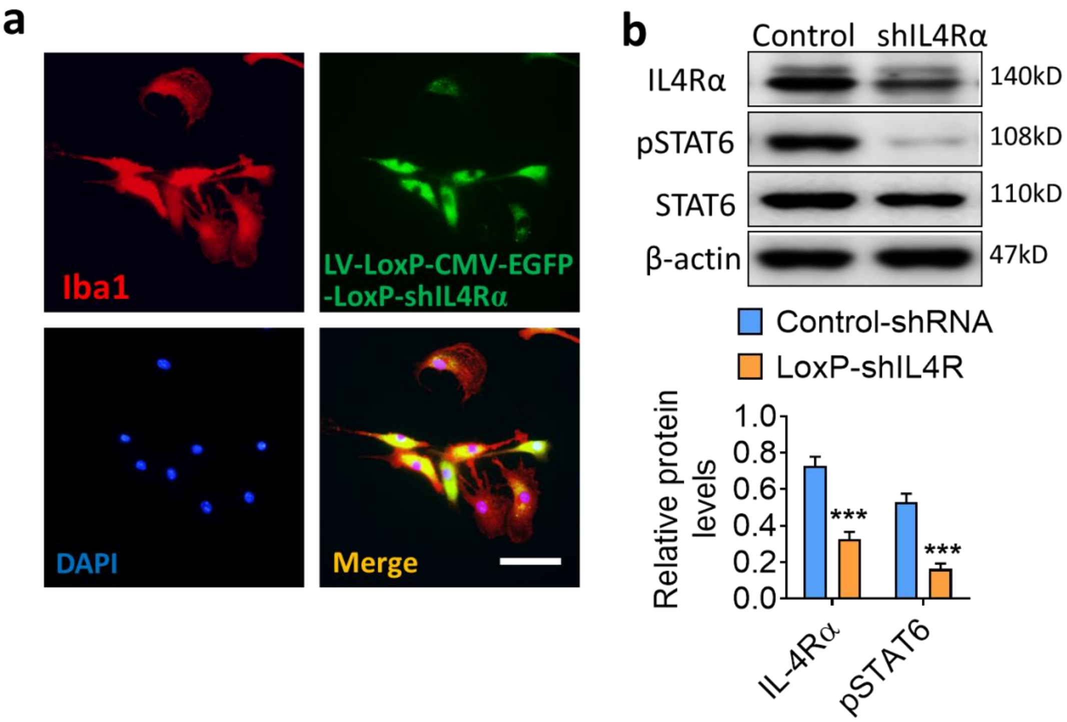


**Fig. S7. Transfection of LV-LoxP-CMV-EGFP-LoxP-shIL4Rα in primary microglia**

**a**, Representative fluorescence micrographs of EGFP, Iba1 and DAPI in primary microglia. Scale bar, 10 μm.

**b**, Western blotting shows the levels of IL4Rα and STAT6 in primary microglia treated by control shRNA or shIL4Rα. IL4 and STAT6 are normalized to β-actin. Data are mean ± SEM (n = 4 per group). *** P < 0.005 (independent-samples t-test).

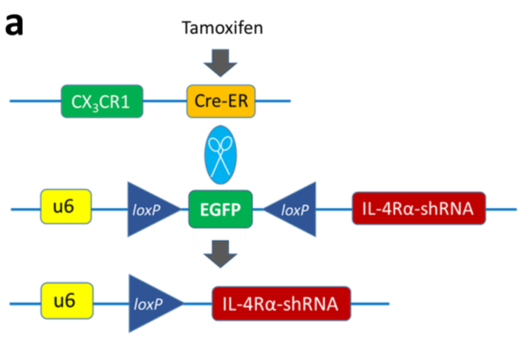


**Fig. S8. Schematic diagram for knockdown of microglial IL4Rα in the hippocampus of $CX_3CR1^{CreER}$ mice**

**a**, Schematic diagram for testing the efficiency of Cre recombinant enzyme in $CX_3CR1^{CreER}$ mice.

**b**, Schematic diagram for knockdown of microglial IL4Rα in hippocampus of $CX_3CR1^{CreER}$ mice.

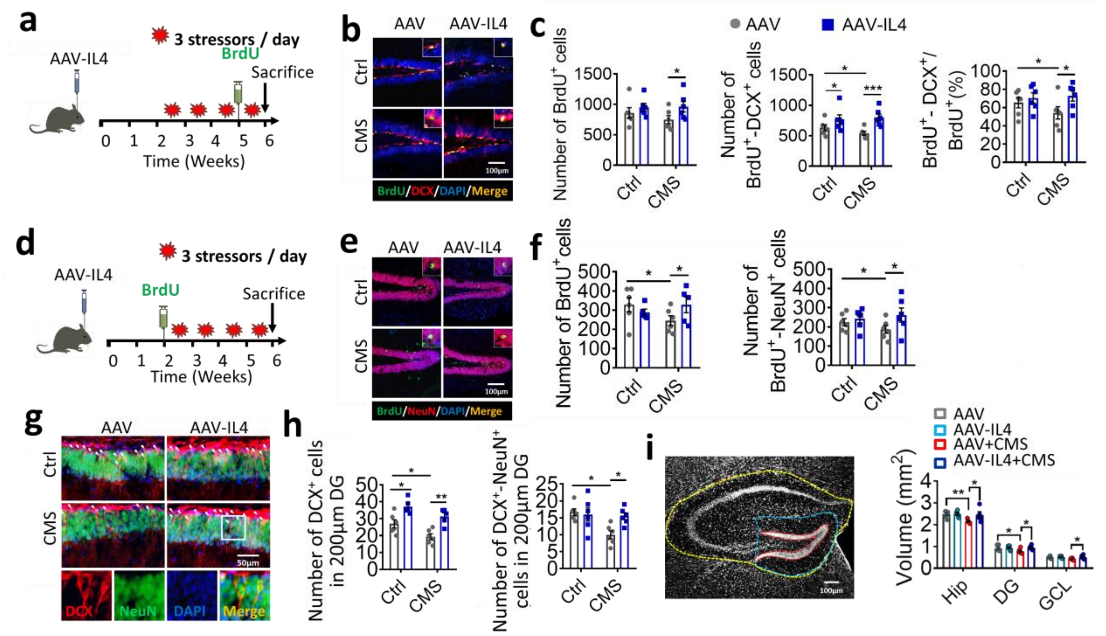


**Fig. S9. Effects of IL4 overexpression on hippocampal neurogenesis in CMS-exposed mice**

**a**, Schematic representation of AAV-IL4 treatment following 4-week CMS, coupled with BrdU injection to monitor proliferating cells within the neurogenic zones.

**b**, Representative fluorescence micrographs showing DCX expression and BrdU incorporation in the SGZ. Scale bar, 100 μm.

**c**, Quantification, per hippocampus, of total number of $BrdU^+$ cells (proliferating cells), total number of $DCX^+$-$BrdU^+$ cells (neuronal progenitors) and percentage of $DCX^+$-$BrdU^+$ cells out of all $BrdU^+$ cells (neuronal differentiation of proliferating cells) in the neurogenic zones.

**d**, Schematic representation of AAV-IL4 treatment following BrdU injection to examine survival and maturity of proliferating cells under stress or without stress.

**e**, Representative fluorescence micrographs showing NeuN expression and BrdU incorporation in the SGZ. Scale bar, 100 μm.

**f**, Quantification of total number of $BrdU^+$ cells and total number of $BrdU^+$-$NeuN^+$ cells per hippocampus.

**g**, Representative fluorescence microscopy images of maturing neuronal progenitors of the hippocampus. Tissue was immunolabelled against DCX and NeuN. Scale bar, 50 μm.

**h**, Quantification of the number of $DCX^+$ cells (per 200 μm of DG) and number of $DCX^+$-$NeuN^+$ cells (per 200 μm of DG).

**i**, Representative micrographs of hippocampus (yellow dotted line), DG (blue dotted line), and GCL (red dotted line), as well as quantification of the volume of hippocampus, DG and GCL. Scale bar, 100 μm. Data are mean ± SEM (n = 5 per group). * $P < 0.05$, ** $P < 0.01$, *** $P < 0.005$ (two-way ANOVA with LSD test).

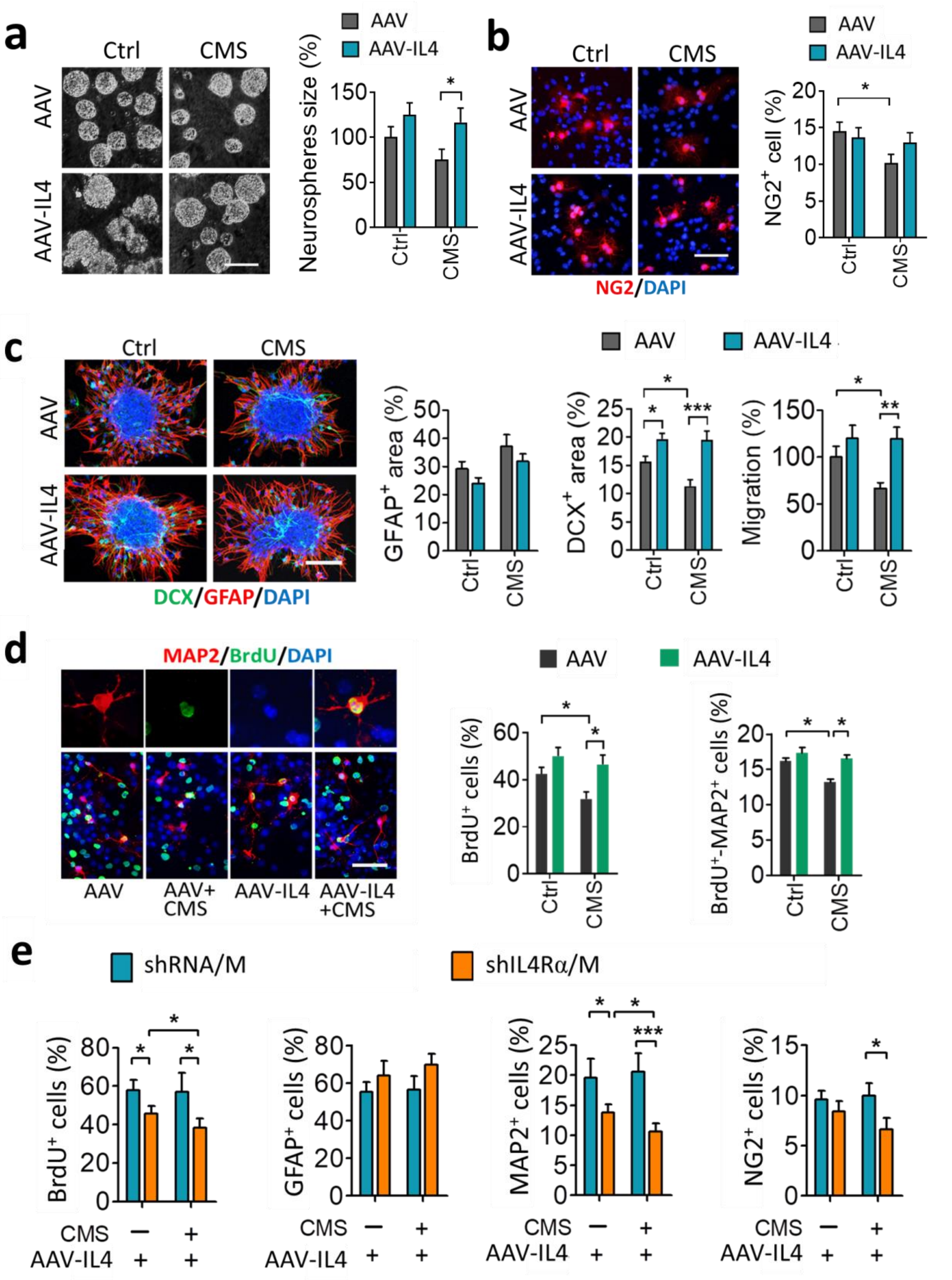


**Fig. S10. Effects of microglia from the hippocampus of AAV or AAV-IL4 mice exposed to CMS on proliferation, differentiation, and survival of NSPCs**

**a**, Effects of microglia on proliferation of neurospheres. Representative micrographs of neurospheres cultured for 3 days in conditioned culture medium from microglia isolated from hippocampus. The neurosphere size are quantified. Data are mean ± SEM (n = 6 per group). Scale bar, 50 μm.

**b**, Effects of microglia on oligodendrogenesis. Representative micrographs of $NG2^+$ cells differentiated from NSPCs when cultured for 3 days in conditioned culture medium from microglia isolated from hippocampus. The number of $NG2^+$ cells is quantified. Scale bar, 20 μm.

**c**, Effects of microglia on differentiation and migration of NSPCs. Representative micrographs of the differentiation and migration of NSPCs cultured for 3 days in conditioned culture medium from microglia isolated from hippocampus. The area of $GFAP^+$ cells and $DCX^+$ cells, and migration of differentiated NSPCs are quantified. Data are mean ± SEM (n = 6 per group). Scale bar, 50 μm.

**d**, Effects of microglia on survival of differentiated NSPCs. Representative fluorescence micrographs of BrdU-labeled NSPCs cultured for 7 days in conditioned culture medium from microglia isolated from hippocampus of AAV, AAV + CMS, AAV-IL4 and AAV-IL4 + CMS mice. Cells were immunolabeled with MAP2 to identify neurons. The percentages of $BrdU^+$ cells (surviving cells) and $BrdU^+$-$MAP2^+$ cells (surviving neurons) are quantified. Scale bar, 20 μm. Data are mean ± SEM (n = 6 per group).

**e**, Effects of microglia in which IL-4Rα was downregulated (before IL-4 overexpression) on NSPCs proliferation and differentiation relative to microglia from AAV-IL4 mice under unstress or stress condition. Data are mean ± SEM (n=5 per group). Data are mean ± SEM (n = 6 per group). * $P < 0.05$, ** $P < 0.01$, *** $P < 0.01$

(two-way ANOVA with LSD test).

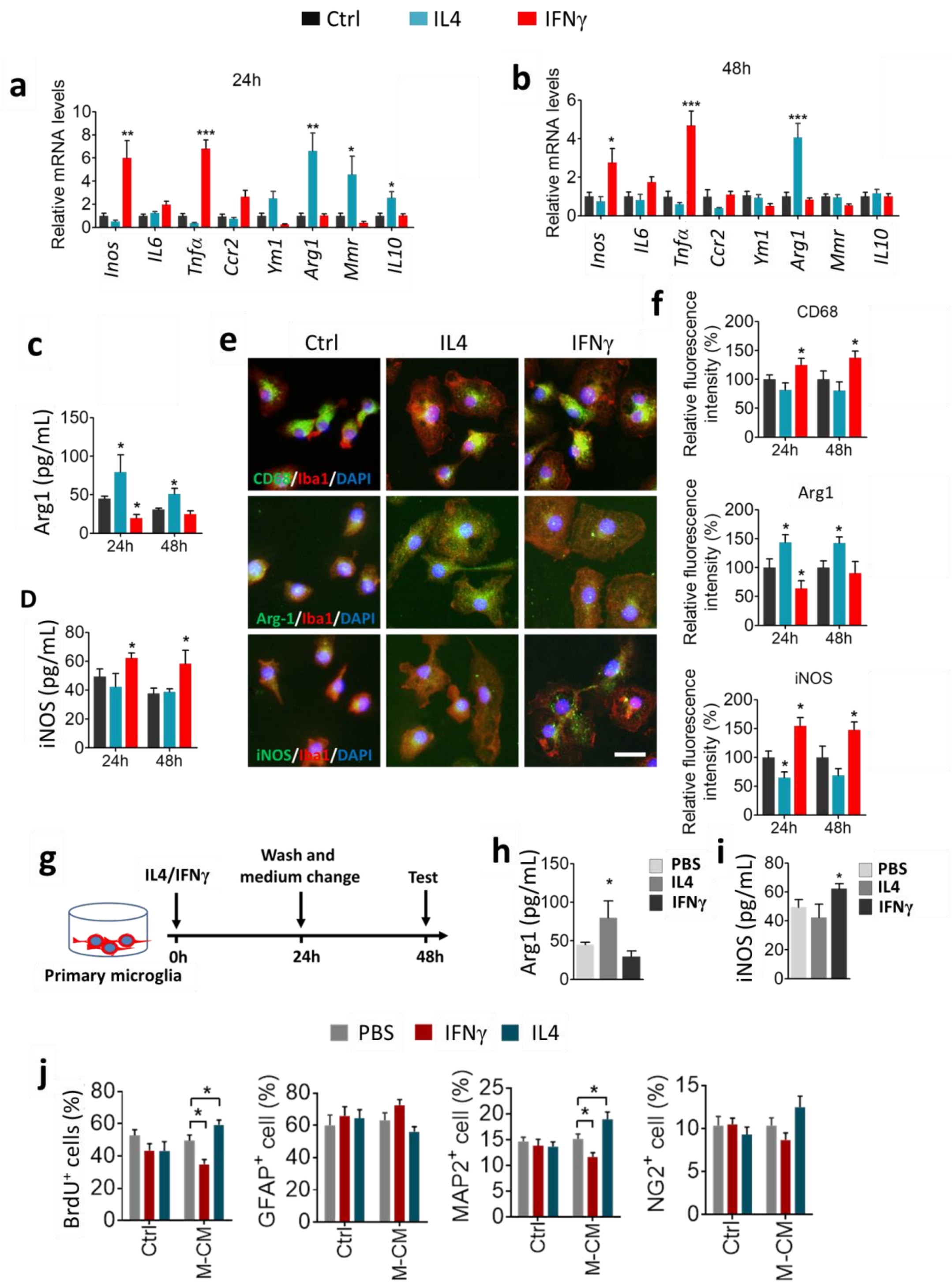


**Fig. S11. The effects of microglia on proliferation and differentiation of NSPCs *in vitro***

**a** and **b**, Changes in mRNA expression of microglial markers after IL4 or IFNγ treatment for 24 h or 48 h.

**c** and **d**, Changes in protein levels of iNOS and Arg1 in microglia for 24 h or 48 h with PBS, IL4 or IFNγ treatment.

**e** and **f**, Changes in fluorescence intensities of CD68, iNOS and Arg1 in microglia for 24 h or 48 h with PBS, IL4 or IFNγ treatment. Scale bar, 10 μm.

**g**, Schematic of the preparation of microglia-conditioned medium (M-CM).

**h** and **i**, Changes in protein levels of iNOS and Arg1 in M-CM. Data are mean ± SEM (n = 4-6 per group). * $P < 0.05$, ** $P < 0.01$, *** $P < 0.005$ (one-way ANOVA with LSD test).

**j**, Effects of PBS, IFNγ, IL4 and M-CM from PBS-, IFNγ-, IL4-treated microglia on proliferation and differentiation of NSPCs. Data are mean ± SEM (n = 4-6 per group). * $P < 0.05$ (two-way ANOVA with LSD test).

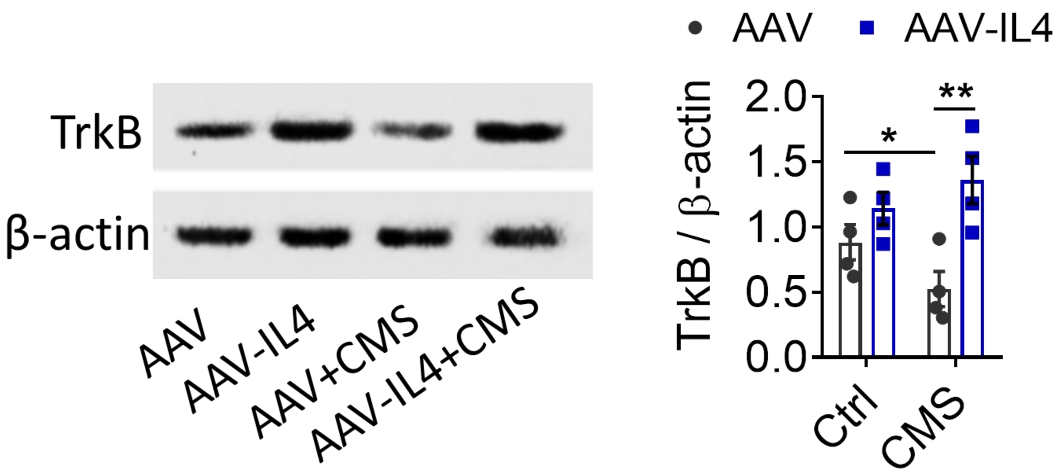


**Fig. S12 Effects of IL4 overexpression on hippocampal TrkB in CMS-exposed mice**

Western blotting shows the levels of TrkB in the hippocampus. Data are mean ± SEM (n = 6 per group). * P < 0.05, ** P < 0.01 (two-way ANOVA with LSD test).

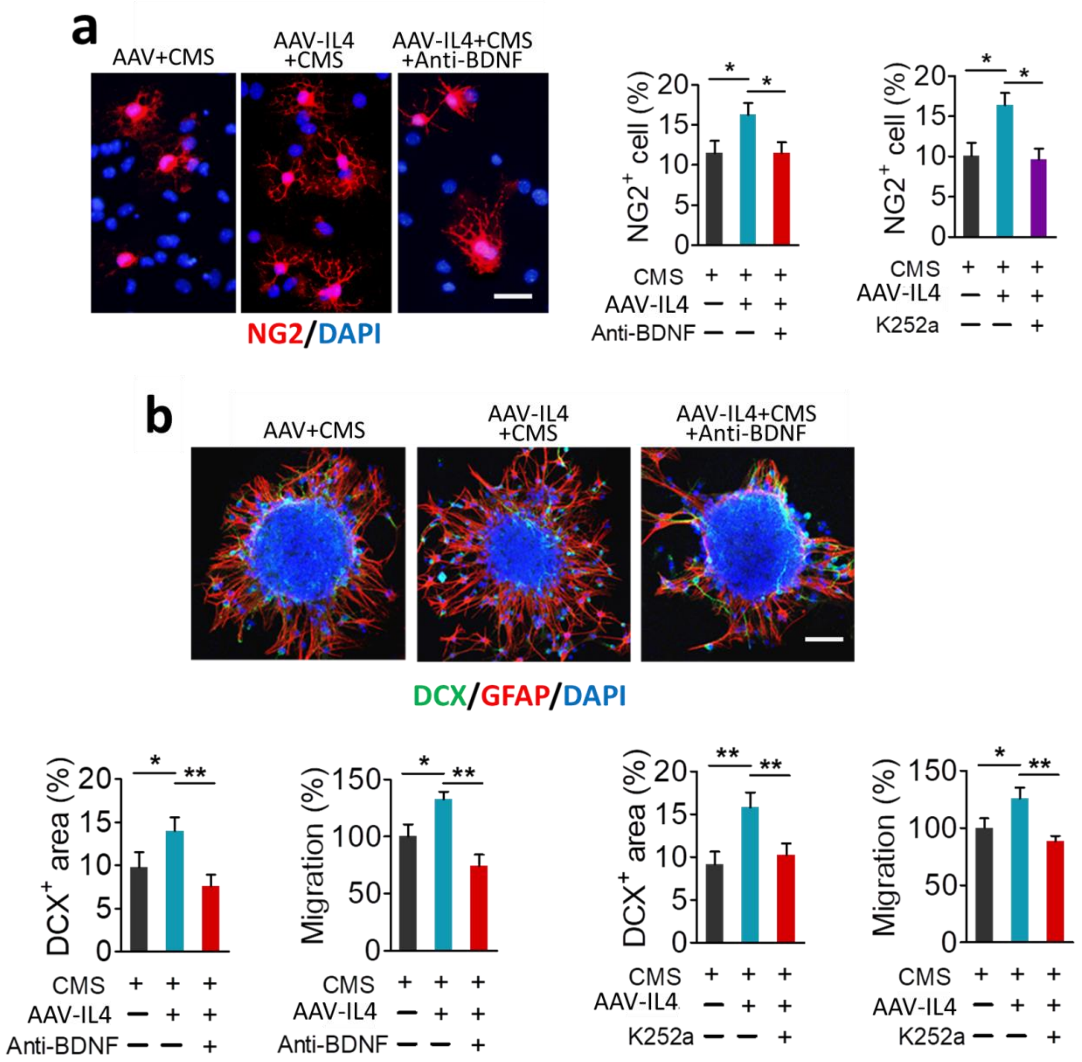


**Fig. S13. Effects of blocking the BDNF pathway on pro-neurogenesis effects of IL4-induced microglia**

**a**, Anti-BDNF antibody or K252a blocked the effects of microglia from hippocampus of AAV-IL4 + CMS on NG2$^+$ cells. Scale bar, 20 μm.

**b**, Anti-BDNF antibody or K252a blocked the effects of microglia from hippocampus of AAV-IL4 + CMS mice on differentiation and migration of NSPCs. Scale bar, 50 μm. Data are mean ± SEM (n = 4–6 per group, 5 replicates per sample). * P < 0.05, ** P < 0.01 (one-way ANOVA with LSD test).

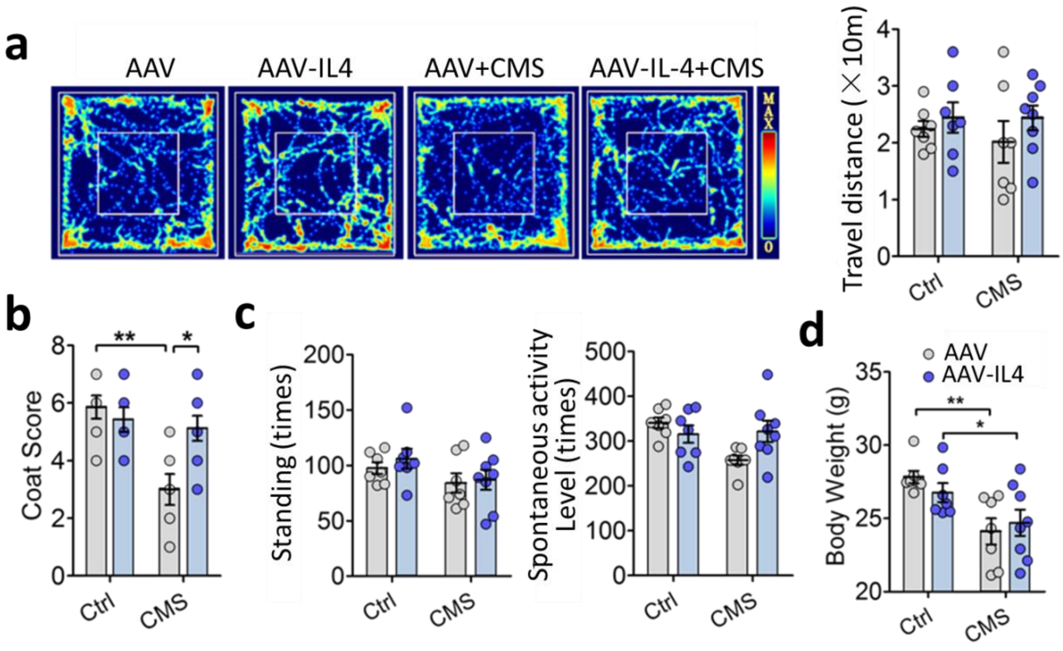


**Fig. S14. Effects of IL4 overexpression in the hippocampus on physiological and behavioral assays of CMS-exposed mice**

**a**, OFT was performed to evaluate travel distance of AAV, AAV + CMS, AAV-IL4 and AAV-IL4 + CMS mice.

**b**, Coat scores were evaluated in AAV, AAV + CMS, AAV-IL4 and AAV-IL4 + CMS mice.

**c**, The number of standing and spontaneous activity levels were evaluated in AAV, AAV + CMS, AAV-IL4 and AAV-IL4 + CMS mice.

**d**, The body weights were evaluated in AAV, AAV + CMS, AAV-IL4 and AAV-IL4 + CMS mice. Data are mean ± SEM (n = 8–10 per group). * $P < 0.05$, ** $P < 0.01$ (two-way ANOVA with LSD test).

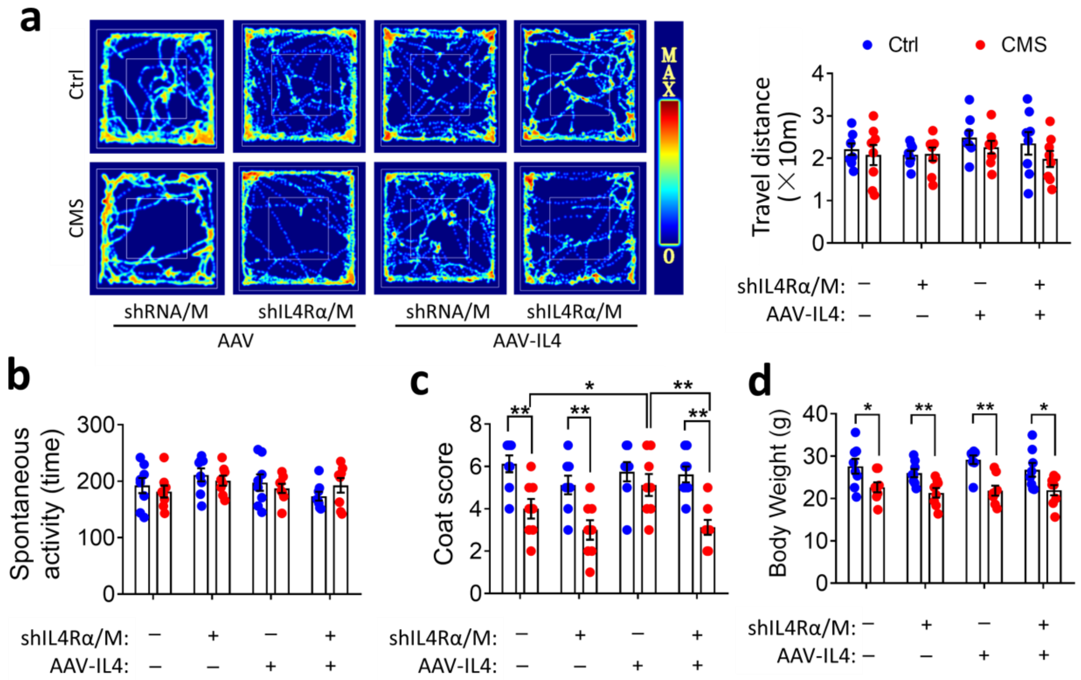


**Fig. S15. Effects of knockdown of microglial IL4Rα on physiological and behavioral assays of AAV-IL4 + CMS mice**

**a**, OFT was performed to evaluate travel distance of all mice.

**b**, The spontaneous activity levels were evaluated in all groups of mice.

**c**, Coat scores were evaluated in all groups of mice.

**d**, The body weights were evaluated in all groups of mice. Data are mean ± SEM (n = 8–10 per group). * $P < 0.05$, ** $P < 0.01$ (two-way ANOVA with LSD test).

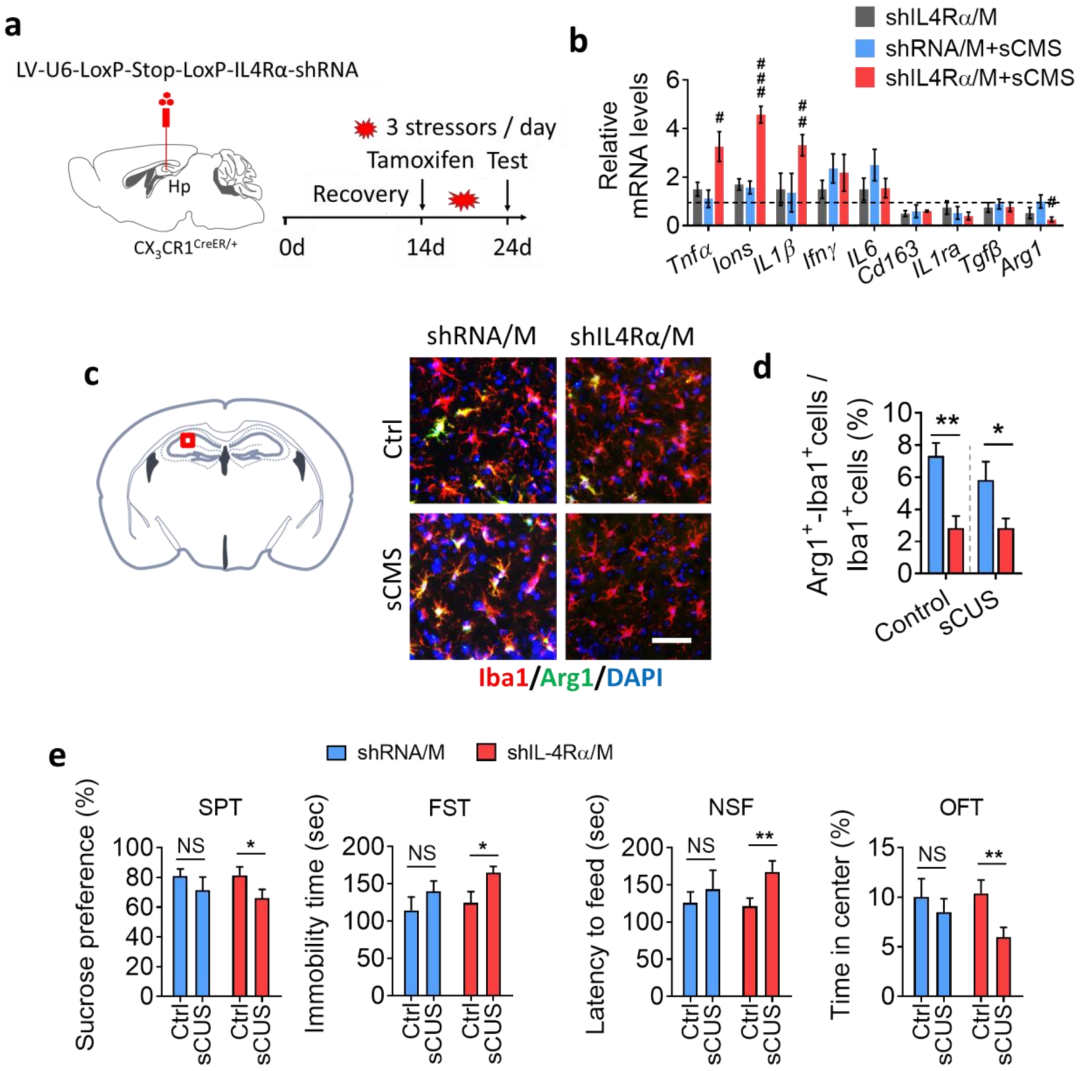


**Fig. S16. Knockdown of microglial IL4Rα increased the vulnerability of mice to stress**

**a**, Experimental timeline of microglial IL4Rα knockdown in the hippocampus and behavioral studies. $CX_3CR1^{CreER}$ mice were injected stereotactically with LV-Loxp-shIL4Rα or LV-Loxp-shRNA into the hippocampus, allowed to recover for 2 weeks, then treated with tamoxifen to induce expression of microglial IL4Rα shRNA. Finally, the animals were subjected to a subthreshold chronic mild stress (sCMS) consisting of exposure to three random stressors daily for 10 days.

**b**, pre-sCMS and knockdown of microglial IL4Rα in the hippocampus significantly increased expression of inflammatory genes. Data are mean ± SEM (n = 4-5 per group). # P < 0.05, ## P < 0.01, ### P < 0.005 *vs* sCMS-shRNA group (two-way ANOVA with LSD test).

**c** and **d**, The percentage of $Arg1^+$ microglia in the hippocampus of mice with pre-sCMS and/or knockdown of microglial IL4Rα treatment. Scale bar, 20 μm. Data are mean ± SEM (n = 5 per group). * P < 0.05, ** P < 0.01 (two-way ANOVA with LSD test).

**e**, Pre-sCMS and downregulation of microglial IL4Rα expression in the hippocampus induced depressive-like behaviors, as assessed by SPT, FST, NSF and OFT. Data are mean ± SEM (n = 9-10 per group). * P < 0.05, ** P < 0.01 (two-way ANOVA with LSD test).

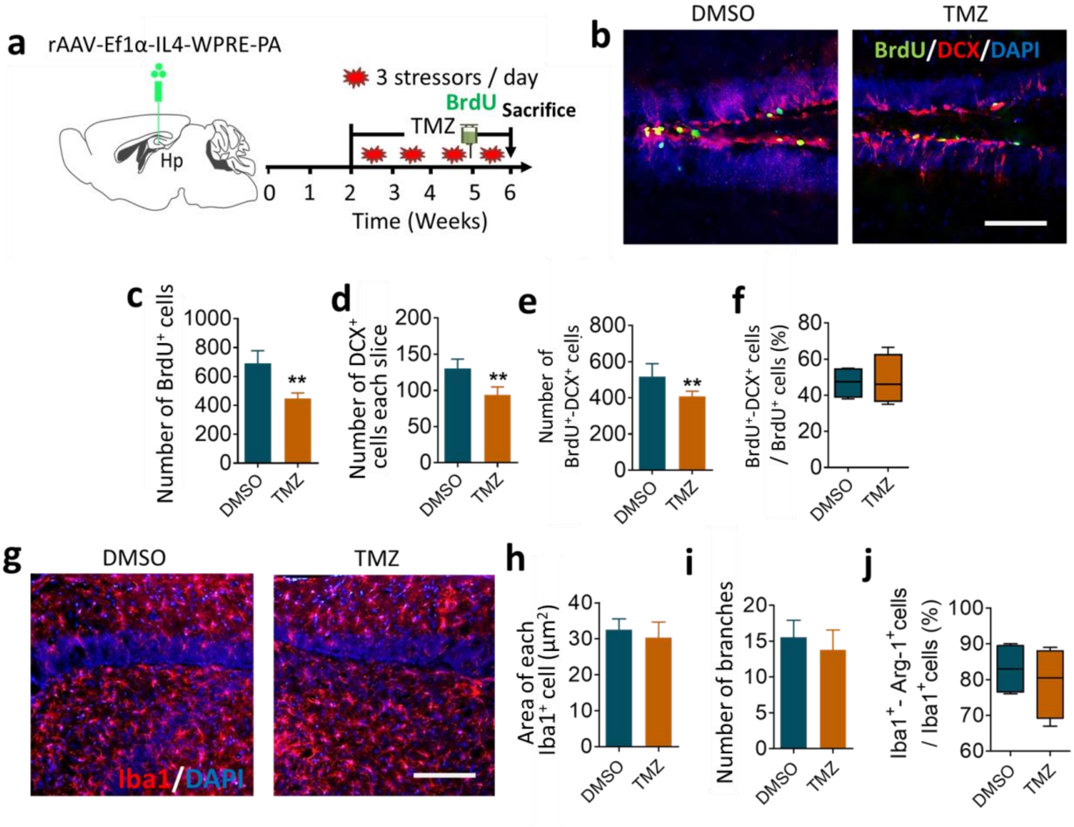


**Fig. S17. Effects of TMZ treatment on hippocampal neurogenesis and microglial morphology**

**a**, Schematic representation of TMZ treatment during 4-week CMS, coupled with BrdU injection to monitor proliferating cells within the neurogenic zones of IL4 mice.

**b**, Representative fluorescence micrographs showing differentiation of proliferating cells in the hippocampus of AAV-IL4-CMS + DMSO or AAV-IL4-CMS + TMZ mice. Scale bar, 100 μm.

**c**, Quantification of BrdU$^+$ cells per DG of mice. Data are mean ± SEM (n = 6 per group).

**d**, Quantification of DCX$^+$ cells per slice in AAV-IL4-CMS + DMSO or AAV-IL4-CMS + TMZ mice. Data are mean ± SEM (n = 6 per group).

**e**, Quantification of BrdU$^+$-DCX$^+$ cells per DG. Data are mean ± SEM (n = 6 per group).

**f**, The percentage of $BrdU^+$-$DCX^+$ cells out of total $BrdU^+$ cells in AAV-IL4-CMS + DMSO or AAV-IL4-CMS + TMZ mice. Data are mean ± SEM (n = 6 per group).

**g**, Representative fluorescence micrographs of microglia in the hippocampus of AAV-IL4-CMS + DMSO or AAV-IL4-CMS + TMZ mice. Scale bar, 100 μm.

**h-j**, Quantification of $Iba1^+$ cells, area of Iba1 positive staining and percentage of $Iba1^+$-$Arg1^+$ cell out of total $Iba1^+$ cells in the hippocampus of AAV-IL4-CMS + DMSO or AAV-IL4-CMS + TMZ mice. Data are mean ± SEM (n = 6 per group). ** $P < 0.01$ (unpaired *t* test)
